# FFPERescuer: deep unsupervised domain adaptation for the reconstruction of gene expression profiles derived from formalin-fixed paraffin-embedded samples

**DOI:** 10.64898/2026.09.02.748790

**Authors:** Lingli He, Kai Song, Ying Li, Yujuan Dong, Cherry Y N Wong, Lin Qi, Xianrui Zhang, Kristiaan Lenos, Tim de Back, Clara Elbers, Chenchen Xu, Royce Man Hin Leung, Renfu Deng, Yinghan Zhang, Sitan Qiao, Feng Gao, Yufeng Chen, Simon Siu-Man Ng, Shentao Zhou, Louis Vermeulen, Xin Wang

## Abstract

Formalin-fixed paraffin-embedded (FFPE) tumor tissues often suffer from RNA degradation, posing a long-standing challenge for reliable transcriptomic profiling. Here, we propose FFPERescuer, a deep learning framework employing unsupervised domain adaptation, to rectify distorted gene expression data. FFPERescuer comprises a partial encoder that maps a small subset of genes to high-level representations and a decoder to reconstruct full gene expression profiles. On simulated data with varying noise levels, FFPERescuer faithfully recovered gene expression profiles, achieving high Pearson correlation coefficients (PCCs > 0.85) with the ground truth. In FF-FFPE-matched cohorts, FFPERescuer significantly enhanced expression profile concordance, with average PCCs increased by 23% (*P* < 0.05). Applying to cancer subtyping, FFPERescuer improved classification accuracy from 67% to 92%, recapitulated subtype-specific biological properties lost in the FFPE-derived data, and enhanced survival associations. Our studies provide a powerful framework for reliable transcriptomic profiling from FFPE-archived tumor samples that are widely available in the clinic.

## 1 Introduction

Next-generation sequencing (NGS)-based transcriptomic profiling of cancer tissues has greatly advanced our understanding of cancer progression, diagnosis, and treatment^1–3^. In practice, fresh frozen (FF) tissue samples are preferred for RNA sequencing (RNA-seq) because they better preserve RNA integrity and generally yield high-quality transcriptomic data. However, FF tissues are not easily accessible as the collection and storage of FF biospecimens are both demanding and costly. In contrast, FFPE tissues are routinely collected in clinical practice, particularly for histopathological diagnosis, and can be stored at room temperature for long periods while largely preserving tissue morphology. Their widespread availability, long-term stability, and rich associated clinical information make FFPE specimens an important resource for studying disease progression, treatment response, and long-term follow-up. A major limitation, however, is that RNA degradation occurs not only during tissue fixation and embedding processes^4^ but also during long-term storage^5^. This degradation could lead to inconsistent results between the matched FF and FFPE samples^6,7^. As a result, how to employ FFPE samples for reliable transcriptomic profiling and downstream analysis remains a long-standing challenge.

Extensive efforts have been made to reduce the influence of RNA degradation on FFPE-derived RNA-seq. On the experimental side, these efforts involve optimizing preanalytical factors during FFPE tissue preparation, including specimen size^4^, ischemic time^8,9^, fixation duration^10^, and storage conditions^11^, selecting RNA extraction protocols^12^, enhancing library preparation methods^13,14^, and optimizing sequencing technologies. For instance, the RNase H method, a type of ribosomal RNA (rRNA) removal method, has been shown to be particularly effective for degraded RNA samples when evaluated based on transcriptome annotation, transcript discovery and gene expression^14^. Recently, NGS technologies specifically tailored for low-quality RNA samples—such as TruSeq (Illumina) and SMARTer (Clontech/Takara Bio)—have also been developed. However, the optimization of these factors has largely been guided by RT-PCR analysis, which offers only a limited view of RNA quality. Consequently, a systematic evaluation of their impact on sequencing data quality at the genome-wide scale remains lacking.

Computationally, the Maxcounts method optimized exon-level expression quantification by leveraging the maximum per-base count, an approach that was robust to biases arising from non-uniform read distribution^15^. The quality surrogate variable analysis (qSVA) employed a structured, data-driven approach to address RNA quality confounding in differential expression analysis^16^. The DegNorm pipeline offers a valuable approach for quantifying both gene- and sample-specific RNA degradation, facilitating its removal as a confounding factor in RNA-seq data analyses^17^. Despite considerable advances, most existing methods are designed primarily to optimize differential analysis outcomes rather than to directly improve the reliability of the underlying gene expression profiles, which are foundational for a wide range of downstream analyses. Furthermore, the application of deep learning approaches specifically aimed at reconstructing reliable FFPE-derived gene expression profiles remain limited.

In recent years, deep learning has emerged as a powerful and general- purpose modeling framework across biomedical domains^18^, highlighted by its capability to automatically learn hierarchical feature representations from raw, high-dimensional data. Unlike traditional statistical models that rely on handcrafted features and strong prior assumptions, deep neural networks can flexibly capture nonlinear relationships, complex dependencies, and context- specific patterns, making them especially well-suited to the inherent complexity of omics data^19^. These general capabilities have led to widespread adoption of deep learning in transcriptomics^20–23^, where gene expression measurements are often noisy, sparse, and confounded by batch effects. By extracting latent representations that reflect underlying biological structure, deep learning models have enabled advances in omics analyses^24,25^ as well as cancer research such as cancer classification^21^, cancer subtyping^20,22^, survival prediction^26^, and drug response prediction^27^.

Parallel to these methodological advances, gene co-expression, defined as the coordinated expression of genes across samples, stands as a well- established principle that fundamentally organizes the transcriptome^28–30^. We hypothesize that genome-wide co-expression patterns present in FF transcriptomic data could be learned and leveraged to guide the reconstruction of FFPE-derived gene expression profiles, and that deep learning is particularly well suited to capture such co-expression patterns and gene-gene dependencies from high-dimensional data. Guided by this rationale, we developed FFPERescuer, a deep learning framework to reconstruct FFPE- derived gene expression profiles from low-quality FFPE inputs. By leveraging gene co-expression patterns^28–30^ and convolutional weight sharing^31^, FFPERescuer aims to narrow the domain gap between FFPE-derived (target domain) and FF-derived (source domain) RNA-seq gene expression profiles.

We evaluated FFPERescuer using both synthetic and real-world datasets, including colorectal cancer (CRC) and high-grade serous ovarian cancer (HGSOC) cohorts, to assess whether it improves concordance between FFPE- derived and FF-derived expression profiles and enhances the utility of FFPE data for downstream biological and clinical analyses. These analyses were designed to evaluate the potential of FFPERescuer as a practical approach for more reliable transcriptomic profiling of archival FFPE specimens.

## 2 Results

### 2.1 TIN is a reliable metric for quantifying RNA integrity in FFPE samples

To systematically quantify RNA degradation, we employed the Transcript Integrity Number (TIN), which measures the uniformity of RNA-seq read coverage across transcripts. High TIN values indicate intact RNA, while low TIN values reflect fragmentation and uneven coverage, leading to quantification biases (**Figure 1A**). Due to RNA degradation, FFPE samples are expected to exhibit substantially lower TINs than matched Fresh-Frozen (FF) samples.

**Figure 1.**
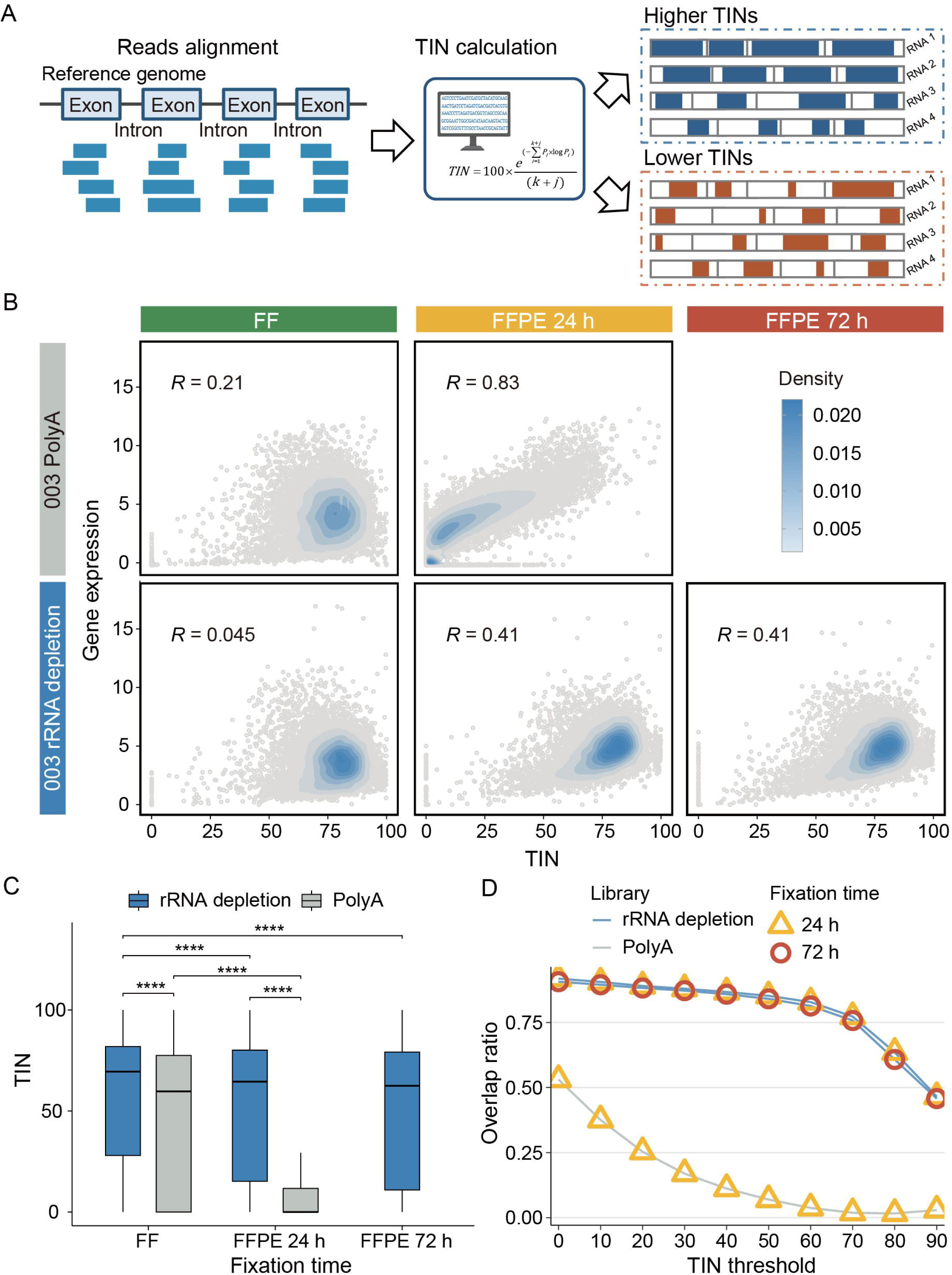
An overview of whole-genome TINs of samples from different experimental settings. **A**, Schematic diagram showing the key points of TIN calculation process. Created in BioRender.com. **B**, Scatter plots showing the correlation between whole-genome TINs and TPM in the whole genome. R values refer to Pearson correlation coefficients. All the correlations are significant. A 2D kernel density estimation of number of points was added to each scatter plot. Darker color indicates larger number of points. **C**, Boxplot depicting whole-genome TINs of the in-house sequenced sample with distinct storage type (FF, FFPE), fixation time (24 h, 72 h), and library preparation strategy (PolyA capture, rRNA depletion). The groups were compared using paired one-sided Student’s t-tests. The center line of the boxplots indicates the median, the edges indicate the interquartile range. **D**, Scatter plot showing the ratio of intersecting genes with TIN above thresholds in FFPE and corresponding FF samples to the total genes with TIN above thresholds in FF samples, under different library preparation methods and fixation durations. All the p-values were Benjamini–Hochberg adjusted. Adjusted *P* ≤ 0.0001 (****).

We first assessed how experimental protocols affect RNA quality using a paired FF-FFPE cohort from the Prince of Wales Hospital (PWH). We evaluated two fixation durations (24h vs. 72h) and two library preparation methods (PolyA capture vs. rRNA depletion) (**Supplementary Table 1**). In FF samples, TIN was largely independent of gene expression level (r = 0.045–0.21, *P* < 0.001). In contrast, FFPE samples showed a strong positive correlation (r = 0.41–0.83, *P* < 0.001, as degradation disproportionately affected lowly expressed genes (**Figure 1B**). We found that longer fixation times significantly reduced TINs, and PolyA-based libraries were markedly more sensitive to fixation duration than rRNA depletion-based libraries (*P* < 0.001, **Figure 1C, Supplementary Figure 1**). The overlap ratio of highly intact genes, i.e., genes with high TINs, between FF and FFPE samples dropped sharply for PolyA-based libraries but remained stable for rRNA depletion-based libraries across increasing TIN thresholds (**Figure 1D**). Crucially, a core subset of genes consistently exhibited high TIN values across all conditions, providing a strong foundation for reconstructing the full transcriptome based on gene co-expression patterns.

To quantitatively assess RNA quality in sequencing data, we calculated TIN for paired FFPE and FF tumor tissues samples from the Prince of Wales Hospital (PWH) cohort, encompassing samples with two fixation durations (24 h and 72 h) and two library preparation methods (PolyA capture and rRNA depletion) (**Supplementary Table 1**). The rRNA depletion library construction method refers to removing rRNA from total RNA. In FF-derived samples, weak correlations were observed between TIN and gene expression levels (R = 0.045–0.21, **Figure 1B**), as high TIN values could correspond to either highly expressed or lowly expressed genes. In contrast, due to RNA degradation, more low TIN values emerge in FFPE-derived samples, resulting higher correlations (R = 0.41–0.83, **Figure 1B**). Further comparisons of TIN values within the paired samples revealed that TINs decreased significantly with longer fixation time and that PolyA-based sequencing data showed significantly lower TINs than rRNA depletion-based sequencing data (all *P* < 0.001, **Figure 1C**; **Supplementary Figure 1**). Furthermore, the degree of decline in TIN for the PolyA-based sequencing data was evidently faster than that for the rRNA depletion-based sequencing data (**Figure 1C**). These results indicate that a sample fixation time of 24 h is more appropriate, and the polyA-based library preparation shows greater sensitivity to fixation duration, while the rRNA depletion-based approach maintains stable TIN values across different fixation durations.

To systematically investigate the impact of fixation time and library preparation strategies, we calculated the overlap ratio of FF samples and FFPE samples with distinct experimental conditions. The results showed that the overlap ratio for the PolyA-based library preparation approach dropped sharply as the threshold increased. In contrast, for the rRNA depletion-based method, whether at 24 or 72 hours of fixation, a noticeable decline in the overlap ratio was only observed when the threshold exceeded 70 (**Figure 1D**). Additionally, regardless of the library preparation method or TIN threshold used, a subset of genes exhibiting uniformly high TIN values was consistently identified in the paired samples. This observation, combined with the well-established principles of gene co-expression pattern, provides a plausible rationale for reconstructing full transcriptome using a small subset of genes.

### 2.2 A genome-wide landscape of TIN in FFPE samples from six types of cancer

Building upon the general analysis of multifactorial influences on RNA quality, this section provides a systematic characterization of the effects of RNA degradation on RNA integrity. Using TIN as a metric, we assessed the RNA integrity of FFPE-sequenced samples from six cancer types. Based on an analysis of 628 FFPE tumor samples, we found that the TIN means for most genes were below 20 across the six cancer types at both chromosomal (**Figure 2A; Supplementary Figure 2**) and gene levels (**Figure 2B**). We further validated the consistency of low TINs across the six cancer types by correlation analysis (all *P* < 0.001, **Figure 2C**). To further confirm low TINs in FFPE samples, we collected 145 FF-derived sequencing samples as an example for comparison analysis. Chromosomal-level TIN means were significantly higher in FF CRC samples compared to FFPE CRC specimens (**Figure 2A; Supplementary Figure 2**). For a given gene, the TIN mean value from the FF CRC (outermost circle) was considerably higher than that from the FFPE CRC (outermost second circle) (**Figure 2B**). These results indicate remarkably lower RNA integrity in FFPE samples compared to FF samples. To quantitatively demonstrate these differences, statistical analyses of genome-wide and sample-wide TIN differences between the CRC FF and CRC FFPE samples were performed. To ensure comparability of TIN values, we restricted the analysis to FFPE CRC samples that used the same library preparation protocol (total RNA-based) as the 145 FF CRC samples. Substantial differences in mean TIN values were observed between the two sample sets (**Supplementary Figure 3**).

**Figure 2.**
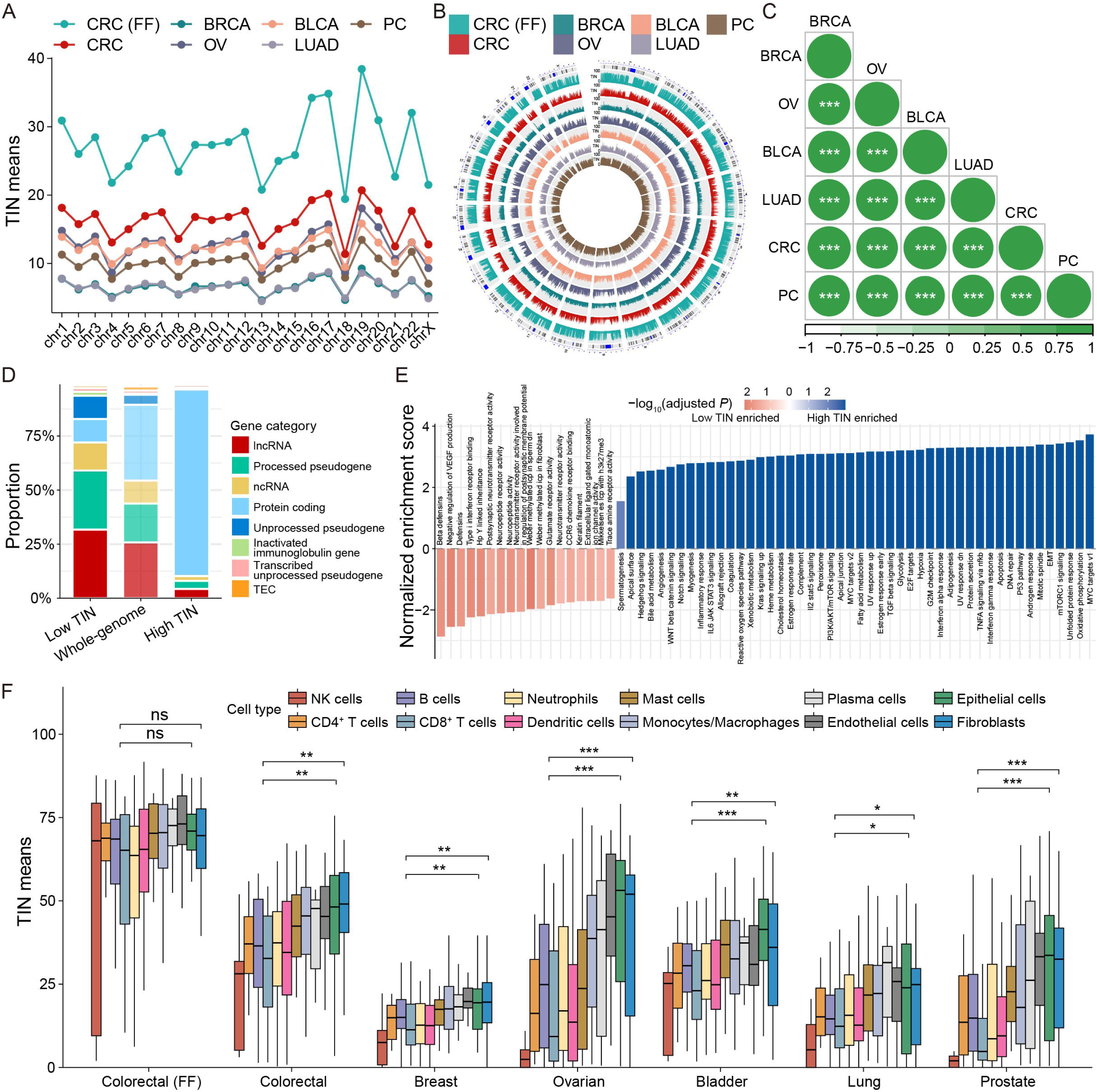
Genome-wide landscape of TINs in FFPE tumor samples from six cancer types. A, Line chart depicting the TIN means within each chromosome across six types of FFPE-derived tumors and one CRC FF-derived samples. **B**, Circos plot showing the whole-genome TIN means for six cancer types with FFPE samples and one cancer type with FF samples. Each circle, colored distinctly, represents a specific cancer type, with the TIN value for each gene indicating the mean TIN within that cancer type. **C,** Correlogram showing the Pearson correlation of whole-genome TIN means between different types of cancer. **D**, Bar plot showing RNA biotype distributions in the gene set with consistently low and high TIN values across six cancer types and genes in the whole-genome. The p-value was determined with Pearson’s chi-square test. **E**, Enriched pathways of the and high TIN gene sets derived from GSEA. **F,** Boxplot showing the TIN means of microenvironment signatures in multiple cancer samples. The statistical difference between NK cells, B cells, CD4^+^ T cells, CD8^+^ T cells, Neutrophils, and Dendritic cells signatures versus epithelial and fibroblast cell signatures were derived from the unpaired one-sided Wilcoxon signed-rank tests. The center line of the boxplots indicates the median, the edges indicate the interquartile range. All the p-values in this figure were Benjamini–Hochberg adjusted. Adjusted *P* ≤ 0.001 (***); Adjusted *P* ≤ 0.01 (**); Adjusted *P* ≤ 0.05 (*); Adjusted *P* > 0.05 (ns: not significant).

To systematically investigate genes with distinct susceptibility to RNA degradation, we compared the biotype composition and functional characteristics of low- and high-TIN genes. Low-TIN genes were enriched for non-coding transcripts, comprising 31.7% lncRNAs, 39.4% processed and unprocessed pseudogenes, and 13.3% ncRNAs, whereas only 9.3% were protein-coding genes. In contrast, high-TIN genes were predominantly protein- coding (86.8%), with much lower proportions of lncRNAs (5.1%), pseudogenes (3.6%), and ncRNAs (1.9%). The whole-genome background showed an intermediate composition, with 25.8% lncRNAs, 22.6% pseudogenes, 10.6% ncRNAs, and 35.0% protein-coding genes (**Figure 2D**). Consistently, lncRNAs were significantly overrepresented in low-TIN genes compared with high-TIN genes (31.7% vs 5.1%, *P* < 0.001).

Gene set enrichment analysis further revealed distinct functional biases between low- and high-TIN genes. Low-TIN genes were preferentially enriched in immune defense and other microenvironment-related pathways, whereas high-TIN genes were enriched in multiple oncogenic pathways (**Figure 2E; Supplementary Tables 2-3**). To further validate the preferential vulnerability of immune-related genes, we examined TIN values across cellular microenvironment signatures derived from pan-cancer single-cell analysis^32^. In FFPE samples, immune signature genes, including NK cells, B cells, CD4^+^ T cells, CD8^+^ T cells, neutrophils, and dendritic cells, showed significantly lower TIN values than epithelial and fibroblast signature genes (**Figure 2F**). In contrast, no significant differences were observed among these signatures in FF samples (**Figure 2F**). Moreover, non-cell-type-specific genes, assessed using 50 GSEA hallmark gene sets, showed no notable TIN variation (**Supplementary Figure 4**). Together, these findings suggest that RNA degradation in FFPE samples may preferentially affect non-coding and immune-related genes.

### 2.3 Unsupervised domain adaptation for reconstructing gene expression profile from FFPE-derived RNA-seq

To resolve the critical issue of poor RNA quality in FFPE samples, we developed a deep learning network to computationally rectify the distorted gene expression data. Our reconstruction framework draws conceptual inspiration from the Speech2face network^33^, adapting its encoder-decoder paradigm where an audio encoder couples with a pretrained image decoder for voice-to- face synthesis, to instead bridge FFPE and FF RNA-seq gene expression profile domains. Specifically, we first constructed a convolutional autoencoder network to learn the co-expression pattern of most of the genes from FF TCGA pan-cancer samples. In this network, the encoder receives the FF gene expression profile as input to extract high-level features, while the decoder reconstructs the gene profile from the extracted features. These modules are referred to as FF-decoder and FF-encoder, respectively.

Second, as the autoencoder network was trained on the lossless FF samples, we subsequently trained a separate partial FF-encoder, for application to the distorted FFPE-derived gene expression profiles. Partial FF- encoder employed the same samples used in training the autoencoder. The distorted expression of whole-genome genes due to RNA degradation as an input can substantially affect the training of the network. Additionally, a large number of redundant genes existed in the FF-derived gene expression profile. To address these challenges, we selected a customized small set of genes, from the FF-encoder input, as an input. Partial FF-encoder was optimized by minimizing the feature differences between itself and FF-encoder, enabling it to extract representative features from gene expression profiles with a small set of genes in a manner as the FF-decoder extracted from FF-derived gene expression profiles. Finally, we combined the partial FF-encoder with the FF- decoder to build the complete reconstruction architecture, designated as FFPERescuer (**Figure 3**). This network performs unsupervised domain adaptation to enable accurate inference of FFPE gene expression profiles from FF gene expression profiles.

**Figure 3.**
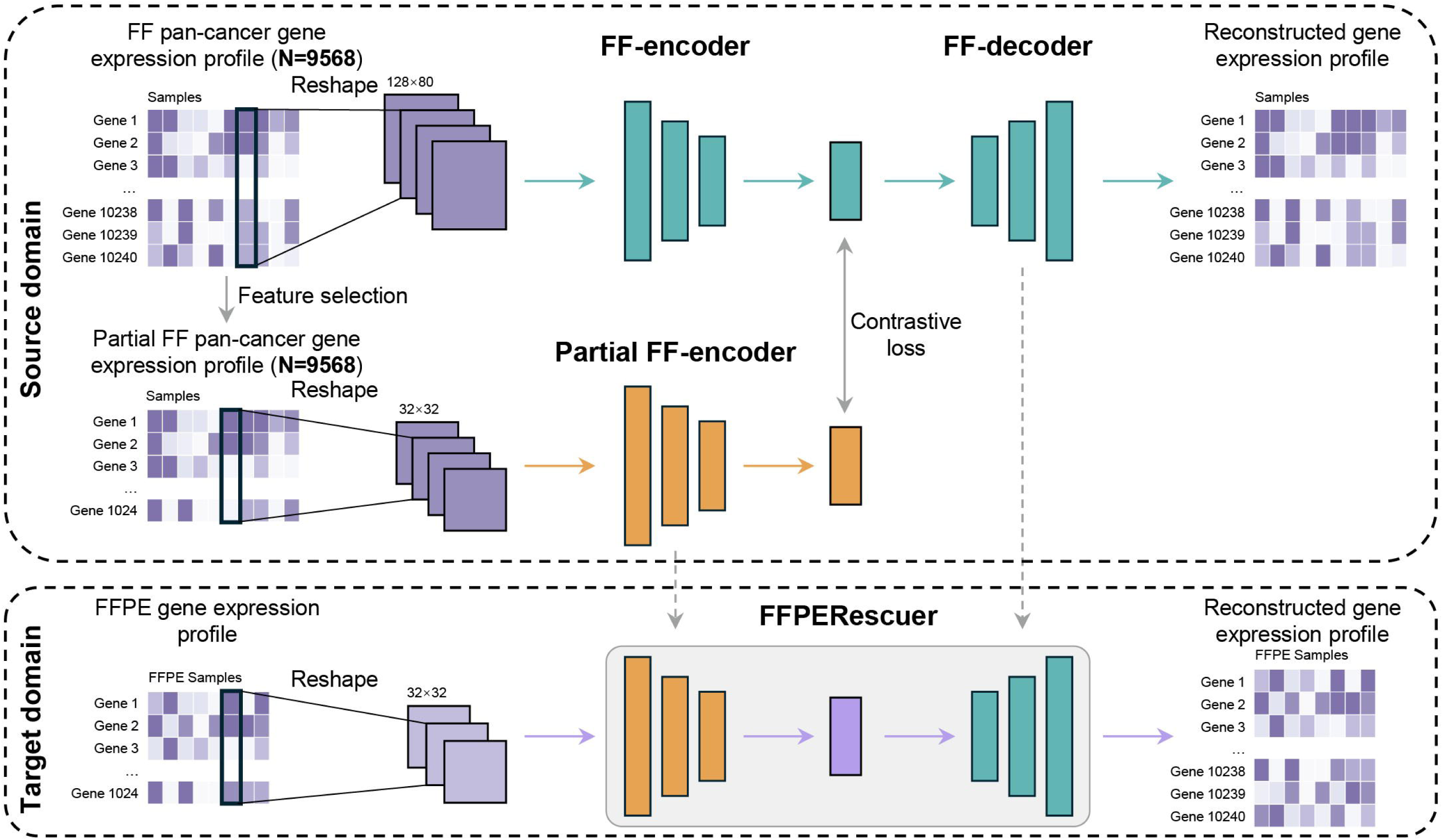
Unsupervised domain adaptation for FFPE-derived RNA-seq profile reconstruction. Schematic diagram of reconstructing FFPE-derived gene expression profile including training two networks and combining them. In the first step, the pan-cancer gene expression profile from FF tissue (source domain) in TCGA was used to train an CNN-based autoencoder network (comprising FF-encoder and FF-decoder). Then, a part the FF pan-cancer gene expression profile with fewer number of genes was used to train the partial FF- encoder. In the third step, partial FF-encoder and FF-decoder were combined to create FFPERescuer, amis for reconstructing FFPE-derived gene expression profile (target domain). FFPERescuer takes a small number of input genes while generates high-dimensional gene expression outputs. CNN, convolutional neural network.

We trained FF-encoder and FF-decoder using the FF TCGA pan-cancer data. Taking the AMC-FFPE CRC cohort as an example, the partial FF-encoder was trained on TCGA FF data using gene features selected from the FFPE cohort. To evaluate the training performance of FFPERescuer, we visualized the features extracted by both networks using t-distributed stochastic neighbor embedding analysis (**Figures 4A–B**). The features derived from both encoders clearly distinguished tumor tissue types within the FF samples, indicating that the networks successfully captured relevant biological characteristics.

**Figure 4.**
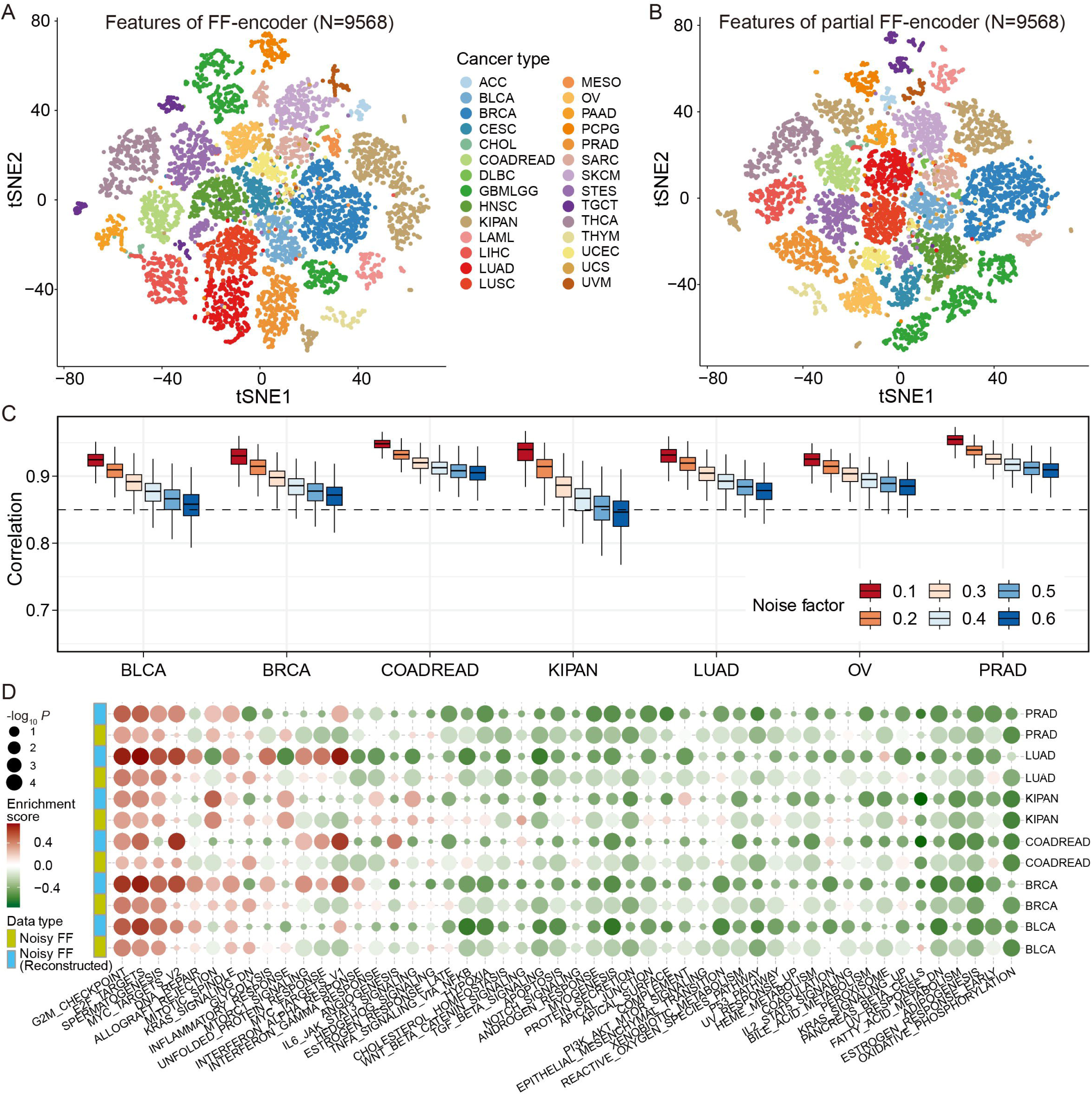
Training performance of FFPERescuer and its reconstruction effectiveness on simulated FFPE cancer samples. **A–B**, t-distributed stochastic neighbor embedding (t-SNE) plots depicting features extracted from the FF-encoder **(A)** and the Partial FF-encoder **(B)**. Each dot denotes one FF sample. Different colors represent distinct types of cancer. **C**, Boxplot showing the Pearson correlations between reconstructed noisy gene expression and FF gene expression in paired samples from seven cancer types. Six sets of noisy gene expression profiles were generated with increasing noise (noise factor equals to 0.1, 0.2, 0.3, 0.4, 0.5, and 0.6). Black dashed line, *R* = 0.85. **D**, Dot plot showing the enrichment scores of GSEA derived from cancer samples versus normal samples. Tumor gene expression profiles were from noisy samples and reconstructed noisy samples respectively. P*-*values were computed using moderated t-tests and adjusted by Benjamini–Hochberg method with R package limma. Noise factor = 0.4.

We then evaluated the robustness of FFPERescuer with simulation data by adding noise to FF samples (see **Methods**). Due to the limited FFPE samples of different cancer types, we did not conduct simulation analysis on all cancer types. For each level of added noise, we computed the sample-wise correlations between both (i) the original and the unreconstructed noisy expression profiles, and (ii) the original and the reconstructed profiles. The correlations between the original and unreconstructed noisy profiles decreased sharply with increasing noise intensity (**Supplementary Figure 5**). In contrast, correlations between original and reconstructed profiles remained consistently high (mostly above 0.85) despite elevated noise levels (**Figure 4C**), demonstrating the robustness of the network-based reconstruction in preserving biological signals.

To assess the biological relevance of FFPERescuer, we applied it under a noise factor of 0.4 to evaluate whether reconstructed gene expression profiles were more informative than their unreconstructed counterparts. GSEA was performed for each cancer type by comparing noisy FF cancer samples with FF normal samples and by comparing reconstructed noisy FF cancer samples with FF normal samples based on the 50 GSEA hallmark pathways. Interestingly, most pathways were downregulated^34^. Both enrichment scores and statistical significance improved substantially after reconstruction **(Figure 4D**), suggesting that the reconstructed gene expression profiles provide enhanced biological advantage compared to their unreconstructed counterparts.

### 2.4 FFPERescuer effectively reconstructs the gene expression level in real-world FFPE datasets

To further demonstrate the effectiveness of FFPERescuer, we applied FFPERescuer to four real-world FFPE datasets, including three CRC datasets and one high-grade serous ovarian cancer dataset (see **Methods**). Most of the samples from the four cohorts had a TIN median of <50 (**Supplementary Figure 6**), indicating poor RNA integrity. TINs of immune cell signatures were apparently lower than those of signatures of other cell types in the three CRC datasets (**Supplementary Figure 7**); this finding was similar to the results depicted in **Figure 2G**.

We first evaluated the effect of reconstruction on gene expression levels. In the AMC-FFPE CRC, reconstruction significantly enhanced expression of CRC signature genes compared to unreconstructed profiles (**Figure 5A**). The gene expression of the consensus molecular subtype (CMS) signatures^3^ was also remarkably improved following reconstruction in the three CRC datasets: AMC-FFPE CRC, GSE86562^35^, and GSE86564^35^ (**Supplementary Figure 8A**). The whole-genome gene expression level was largely enhanced after reconstruction on AMC-FFPE CRC (**Supplementary Figure 9**). We then calculated Pearson’s correlations between each pair of FFPE and FF samples before and after reconstruction based on 10,240 genes in the three CRC datasets. Pearson’s correlation coefficients between the gene expression of each paired FFPE and FF samples were elevated after reconstruction (**Figures 5B–C**; *P* = 2.4 × 10^-4^, **Figure 5D left**). Significant improvements in correlation were also observed in the other two datasets (*P* = 2.7 × 10^-10^, **Figure 5D middle**; *P* = 2.6 × 10^-10^, **Figure 5D right**). To further quantify the promotion of a single gene’s expression before and after reconstruction, we applied Euclidean distance to particularly determine the reconstruction efficiency of immune cell signatures which showed relatively lower TINs in FFPE samples. The Euclidean distances with the corresponding gene expression in paired FF samples were significantly decreased in the three datasets after reconstruction (*P* = 2.3 × 10^-9^, **Figure. 5E left**; *P* = 0.0083, **Figure 5E middle**; *P* = 3.1 × 10^-5^, **Figure 5E right**). The same analysis for the other cell-specific signatures showed the same reconstruction effect (*P* = 1.8 × 10^-13^, **left**; *P* = 1.1 × 10^-5^, **middle**; *P* = 5.1 × 10^-11^, **right, Supplementary Figure 8B**). To conclude, FFPERescuer can substantially reconstruct the gene expression level for the 10240 genes, particularly for cell-specific signatures in real-world FFPE datasets.

**Figure 5.**
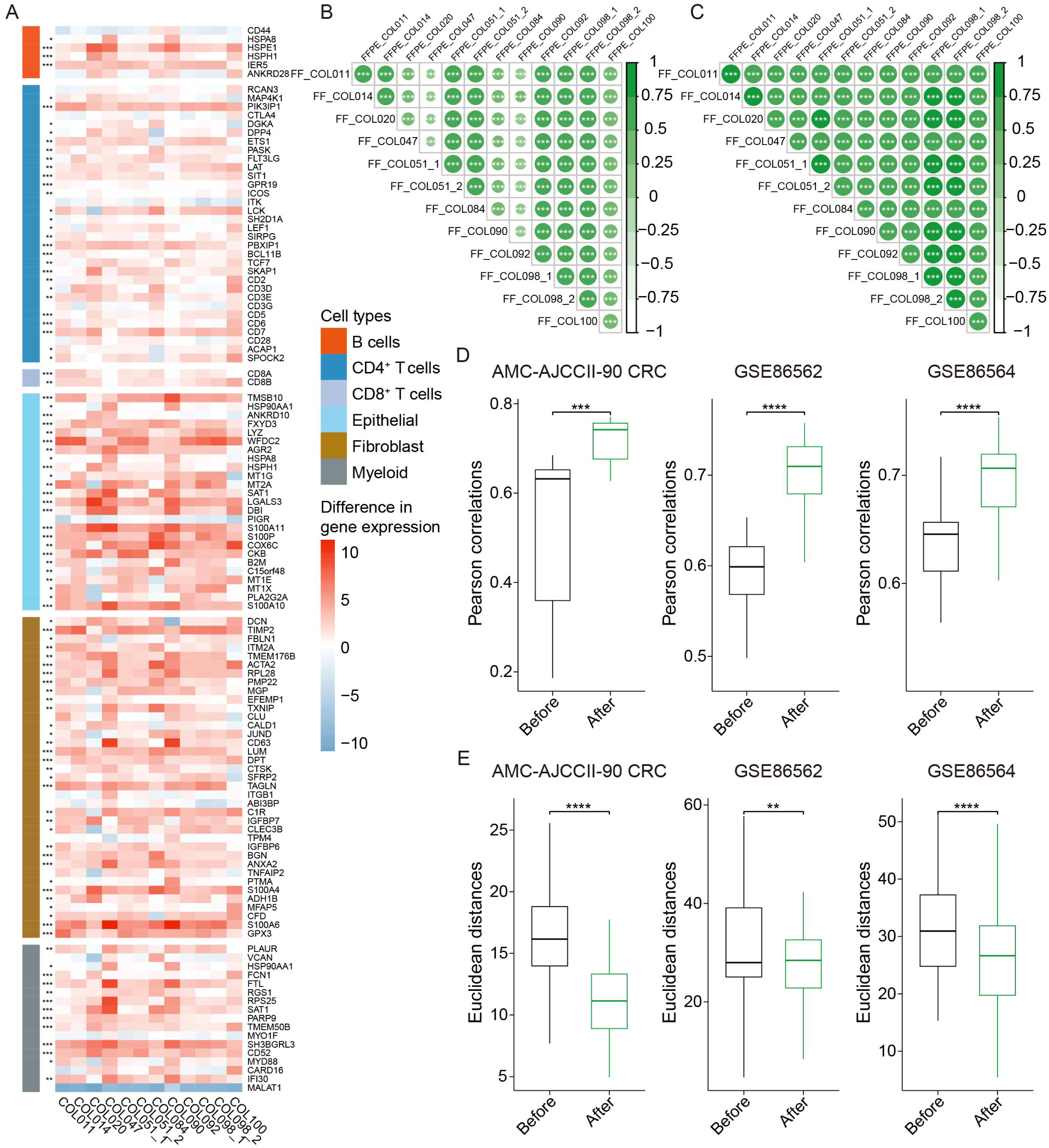
Comparisons of gene expression level of paired real-world CRC FFPE and FF samples before and after applying FFPERescuer. **A**, Heatmap showing the differential expression of cell-specific gene signatures in AMC- FFPE CRC across 12 FFPE samples before and after reconstruction, with significance based on statistical testing of gene expression changes. **B–C**, Correlogram showing the pair-wise sample Pearson correlations between FF expression and FFPE gene expression before reconstruction (**B)** and FFPE gene expression after reconstruction (**C)** respectively on the AMC-FFPE CRC. All sample-wise correlations were calculated based on the 10240 genes. **D**, Boxplots showing the pair-wise sample Pearson correlation between FF gene expression and FFPE gene expression on the three cohorts before and after reconstruction. **E,** Boxplot showing the pair-wise immune cell signature gene Euclidean distances of FF gene expression and FFPE gene expression on the three cohorts before and after reconstruction. The statistical differences were derived from paired one-sided Wilcoxon signed-rank tests for **Figure 5A, 5D**, and **5E**. The center line of the boxplots indicates the median, the edges indicate the interquartile range. All the p-values in this figure were Benjamini–Hochberg adjusted. Adjusted *P* ≤ 0.0001 (****); Adjusted *P* ≤ 0.001 (***); Adjusted *P* ≤ 0.01 (**); Adjusted *P* ≤ 0.05 (*).

### 2.5 FFPERescuer reconstructs faithful gene expression profiles from FFPE samples, enabling accurate molecular subtyping and prognostic stratification

Next, we evaluated the clinical utility of the reconstructed gene expression profiles. Based on the AMC-FFPE CRC cohort, we conducted principal component analysis (PCA) of FFPE gene expression profiles before and after reconstruction using the ground truth of CMS (**Figures 6A–B, Supplementary Table 4**). A large increase in the accumulative variance contribution of PC1 and PC2 (from 69.4% to 88.0%) was observed after reconstruction. The “COL014” sample belonging to CMS4 became closer to the CMS4 group after reconstruction. Following the classification of the samples into before and after reconstruction groups, a large increase in accuracy from 66.67% to 91.67% was observed (**Figure 6C**).

**Figure 6.**
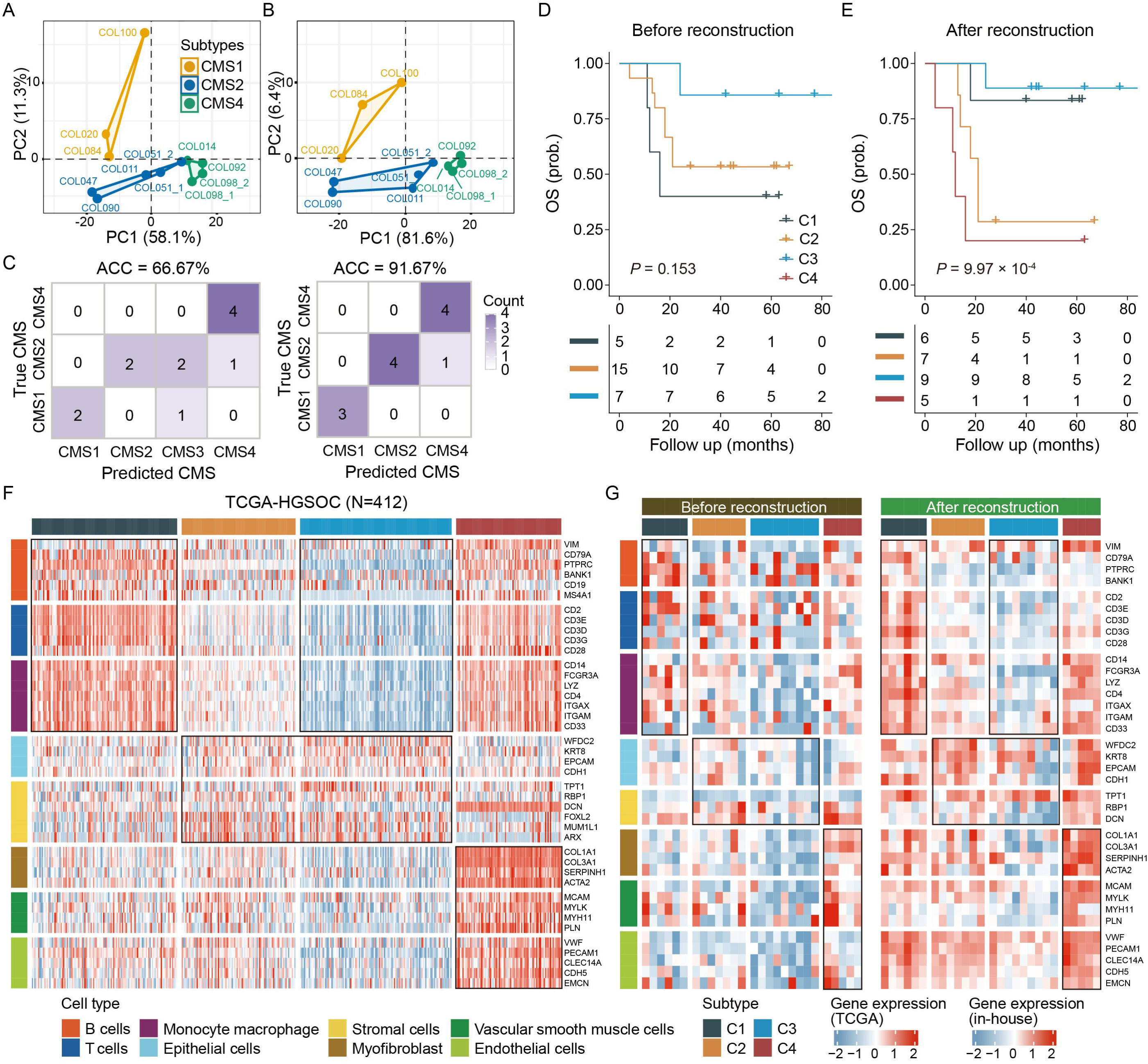
Comparisons of clinical properties between paired FF and FFPE samples (CRC) and unpaired FFPE samples (HGSOC) before and after applying FFPERescuer. **A–B**, PCA plots showing the percentages of variance explained by true CMS before **(A)** and after reconstruction **(B)**. Each dot represents a sample colored with true CMS. **C**, Confusion matrix showing the association between true CMS and predicted CMS results from raw FFPE samples (left) and reconstructed FFPE samples (right). The subtypes were calculated via CMSclassifier. Accuracy: 66.67% (before); 91.67% (after). **D–E**, Kaplan–Meier plots of overall survival for all patients from the West China HGSOC classified before (**D**) and after reconstruction (**E**). The p-values were determined with log-rank tests. **F**, Scaled gene expression of ovarian cancer cell-specific signatures across the TCGA-HGSOC cohort (ordered by subtype assignments). **G**, The same gene order was applied to display expression profiles in the West China HGSOC cohort before and after reconstruction. C1, C2, C3, and C4 denote immunoreactive, differentiated, proliferative, and mesenchymal, respectively.

To demonstrate the broad applicability of our framework to various cancer types, we additionally tested it on a HGSOC FFPE cohort. We applied a four- class HGSOC classifier trained using the subtype information from the study of Konecny *et al*.^36^ (see **Methods**), to obtain molecular subtypes for the FFPE samples. FFPE samples were classified into three subtypes, as the C4- mesenchymal subtype was absent prior to reconstruction, reflecting distortion in the gene expression profile (**Figure 6D**). After reconstruction, the C4- mesenchymal subtype emerged and showed the worst OS (*P* = 9.97 × 10^-4^, **Figure 6E, Supplementary Table 5**). The reconstructed HGSOC cell-specific signatures exhibited expression patterns across subtypes that more closely resembled those observed in TCGA data (**Figure 6F**, **Figure 6G, right**). In contrast, the pre-reconstruction expression profiles failed to demonstrate this concordance (**Figure 6G, left**).

Both quantitative and qualitative evaluations demonstrate that FFPERescuer effectively narrows the domain gap between FFPE and FF- derived gene expression profiles, thereby facilitating more accurate identification of cancer subtypes and biologically relevant features. Furthermore, its successful application to both CRC and HGSOC samples highlights the versatility and broad applicability of FFPERescuer across multiple cancer types.

## 3 Methods

### 3.1 Data collection and curation

The 788 public FFPE pan-cancer samples across 9 cancer types with fastq files were collected from the Gene Expression Omnibus database (GEO, https://www.ncbi.nlm.nih.gov/gds/) (**Supplementary Table 6, Supplementary Figure 10**). After sample filtering (see **Methods**), 628 samples were retained for further analysis. Gene expression profiles of 9568 transcriptomes across 28 types of cancer with FF primary tumor tissues were collected from the TCGA database. Among these, seven cancer types were used for simulation analyses as only these cancer types have FF normal samples in the TCGA database and external FFPE data from 788 FFPE samples (**Supplementary Table 6**). One dataset with 145 fastq files of FF tumor samples was also collected for comparison with real-world data. All specimens were obtained from primary tumors.

The study also included three paired FF and FFPE RNA-seq cohorts and one FFPE-only RNA-seq cohort (**Supplementary Table 7**). All the specimens from the same patients were from the same tumor. The PWH CRC specimens were collected from Prince of Wales Hospital, Hong Kong (**Supplementary Table 1**). The 12 samples of the AMC-FFPE CRC were obtained from the Academic Medical Center, Amsterdam. The matched FF-derived data were deposited in the AMC-AJCCII-90 (GSE33113) dataset. The 27 samples of the West China HGSOC cohort were collected from the West China Second University Hospital, Sichuan University. The public CRC datasets with paired FF (GSE86557^35^) and FFPE (GSE86562, GSE86564) data were obtained from the GEO database. A comprehensive description of the patient cohorts utilized in each analysis can be found in **Supplementary Tables 1 and 4–9**.

### 3.2 Sample sequencing and tumor profiling

For the AMC-FFPE CRC (N=12) cohort, all the total RNAs from FFPE tissues were isolated using the AllPrep DNA/RNA/miRNA Universal Kit (Qiagen, catalog number: 80224) except for two samples (COL051_2, COL098_2), whose total RNAs were extracted with the RNeasy FFPE Kit (Qiagen, catalog number: 73504). TruSeq RNA Exome (Illumina) was employed for library preparation and single-end 65-bp sequencing. The sequencing was performed on the Illumina Hiseq 2500 platform.

For the West China HGSOC cohort, RNA extraction, library construction, and sequencing services were provided by Novogene Co., Ltd. (Tianjin, China). Total RNA was extracted, and libraries were constructed using an mRNA enrichment-based approach. The purified libraries were analyzed by an Illumina Hiseq X Ten platform with 150-bp paired-end reads.

For the PWH CRC cohort, specimens were obtained from three patients with CRC. The recommended fixation duration for RNA is 24–72 h for resection specimens^37^. Therefore, we used 24 and 72 h as our fixation conditions. Total RNA was extracted using the PureLink™ RNA Mini Kit (Thermo Fisher Scientific, catalog number: 12183018A) and the PureLink™ FFPE RNA Isolation Kit (Thermo Fisher Scientific, catalog number: K156002) for FF and FFPE tissues, respectively. The obtained RNAs were then sent to Novogene Co., Ltd. (Hong Kong, China) for quality control. One of the three samples passed the quality control test and were used for library preparation and transcriptome sequencing.

The PolyA library preparation method was the commonly used protocol based on our collected data (**Supplementary Table 6**). The rRNA depletion-based library preparation approach can outperform the PolyA-based library preparation protocol^14^. Therefore, we used these two protocols for library preparation. **Supplementary Table 1** provides detailed information of the RNA-seq for the PWH CRC.

Approvals by ethics committees were obtained at corresponding clinical sites. All patients provided written informed consent.

### 3.3 Data preprocessing

NGS reads were aligned to the Genome Reference Consortium GRCH38 assembly by using “Spliced Transcripts Alignment to a Reference” (STAR, v.2.7.3a)^38^. STAR was conducted with parameters suggested by the ENCODE project. RNA-seq by expectation maximization (RSEM) was used to quantify gene expression^39^ in both FF and FFPE samples. The sorted and indexed BAM files were used to calculate TINs with RSeQC (v 4.0.0)^40^.

The datasets from the different library preparation methods and sequencing methods led to different sequencing results. However, the procedures for TIN calculation do not consider upstream effects. Hence, before merging different FFPE datasets within the same cancer type, we removed batch effects for TINs with the sva package in R software^41^.

We also found that many FFPE samples had very low TINs, with the TIN mean scores almost equal to zero. Because multiple zero values could affect the outcomes of the analyses, we performed sample filtering for the public FFPE samples. Any sample with >75% zero TIN genes were filtered out. We found that most of the discarded samples involved the PolyA-based sequencing method (**Supplementary Figure 11**). This finding was consistent with the conclusions from previous studies and the results of our present study. Finally, 628 public FFPE samples were retained.

### 3.4 Feature selection

Feature selection in FFPERescuer involved two main components corresponding to the input gene sets of the two network modules. For the first component, the log_2_(transcripts per million [TPM]+1) profile was obtained, and the top 10,240 genes with larger median absolute deviation (MAD) values across all 28 cancer types were retained.

The second component involved in partial FF-encoder focused on maintaining feature genes that are both biologically significant and minimally degraded. Since immune-related gene signatures often exhibit low TINs, selecting only high-TIN genes could unintentionally exclude important biological information. To address this, we prioritized cell-type-specific signatures, which may carry crucial information. For the simulation analyses with seven cancer types, the cell-specific signatures derived from pan-cancer single-cell analysis were collected from a published study^32^. If the signature was annotated with unknown or normal enriched in the supplementary table, the genes were excluded from the list. A total of 308 unique signatures from 12 cell types were finally chosen. Among these, genes with a TIN median score of >20 were retained. This can be explained by the positive relationship between gene expression and gene TINs in FFPE samples (**Supplementary Figure 12**). Next, for the excluded signatures (gene set A), we performed Pearson’s correlation analysis between the expression of these genes and all other non-signature genes (gene set B) based on the TCGA FF pan-cancer gene expression data. Any gene in gene set B showing a correlation coefficient of >0.8 with any genes in gene set A and having a TIN median score of >20 was retained. In other words, we aimed to find substitute genes for the signatures with low TINs. Third, to obtain the final list of 1024 genes, we attempted to find genes that had not been included in the input gene list with higher TINs than those of other genes. For the real-world FFPE dataset-based analyses, these three steps were performed, except for signature gene selection in the first step. CRC single cell-specific signatures were mostly obtained from the study of Li *et al*.^42^. T cell-specific signatures from CIBERSORT^43^ were also added to the CRC signature list. A total of 176 unique signatures from 6 cell types were selected. These same signatures were used for the analysis shown in **Figure 4A**. HGSOC single cell-specific signatures were mainly obtained from the study of Zhang *et al*.^44^. T cell-specific signatures from CIBERSORT were also added to the HGSOC signature list. Because different datasets may have different genes with higher TINs (**Supplementary Figure 13**), we trained partial FF-encoder for each dataset individually to make better use of those genes with a more reliable expression pattern.

### 3.5 FFPERescuer architecture and two-stage training strategy

In the first stage, we trained a feature reconstruction autoencoder composed of the FF-encoder and FF-decoder using FF gene expression data. The FF- encoder was implemented as a convolutional neural network (CNN) to extract high-level representation of gene expression. It consisted of three convolutional layers, each followed by a max-pooling layer, and a bottleneck layer implemented via a fully connected layer to capture the compressed feature representation. The FF-decoder mirrored the structure of the encoder and aimed to reconstruct the original gene expression profile from the bottleneck features. Rectified Linear Unit (ReLU) activations were employed throughout to mitigate vanishing gradient issues^45^. The model was optimized by minimizing the mean squared error (MSE) between the input and the reconstructed gene expression profiles. Training was performed using the Adam optimizer^46^ with a learning rate of 1 × 10^-3^. In the second stage, we trained the partial FF-encoder, designed to process distorted FFPE-derived expression profiles. This network shared the same output dimensionality as the bottleneck layer of the FF- encoder. The training objective was to minimize the MSE between the feature output of the partial FF-encoder and the corresponding bottleneck features generated by the FF-encoder, thereby aligning representations across FF and FFPE domains.

We randomly divided the 9,568 FF TCGA RNA-seq transcriptomes into a training set (80%) and a testing set (20%). The deep learning models were implemented in Python using the Keras library (https://keras.io/) with TensorFlow (https://www.tensorflow.org) as the backend.

### 3.6 Data processing of the network output

The input data of FFPERescuer was zero-one scaled. Hence, the output data were not the actual level of gene expression. A reverse scale step was deployed to obtain the final gene expression level comparable to TPM. First, the minimal expression level and the range of expression levels of the 10,240 genes in each cancer type among the 9568 FF transcriptomes were calculated. Second, for a specific type of cancer, the final expression level of each gene was calculated using the equation: *E_reverse_* = *M* + *R* × *E_output_*, where *E_output_* represents the output expression of a given gene, *R* represents the expression level range of the gene in the corresponding cancer type, and M represents the minimal expression level of the gene in the corresponding cancer type.

### 3.7 TIN computation

The Transcript Integrity Number (TIN)^40^ provides a quantitative assessment of RNA integrity by evaluating the uniformity of sequencing coverage across representative transcript positions. The calculation involves: (1) selecting *k* equally spaced nucleotide positions plus all exon-exon junction positions (*j*); (2) computing relative coverage at each position 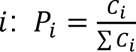, where *C_i_* denotes the read count at position *i* and ∑ *C_i_* represents the total read coverage across all selected positions; (3) calculating Shannon entropy 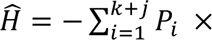 *logP_i_*, which approaches maximum when coverage is perfectly uniform; and (4) deriving the final TIN score:

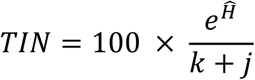

which ranges from 0 to 100, representing the percentage of the transcript with uniform read coverage.

### 3.8 Simulation analysis

To determine the type of simulated noise that should be added to FF samples, we visualized the density distribution of the (100 - TINs) of the network FFPE input genes and found that the scores exhibited approximately normal distribution (**Supplementary Figure 14**). The normal distribution noise of gene expression was used in a previous study^17^. Therefore, we generated random Gaussian noise using different noise factors. Seven cancer types were used for simulations. For each cancer type, the first autoencoder network was the same, and the second network was trained individually. Therefore, each cancer type had its own combined reconstruction network. In our simulations, four distinct levels of noise were first added to the TCGA FF samples 100 times independently. Next, each noisy FF gene expression profile was inputted into the trained FFPERescuer for reconstruction.

### 3.9 Bioinformatics analysis

The low TIN genes were statistically discovered using RankProd R package^47^. The pathway clusters were visualized in cyotscape (v3.10.3). The CMS subtype for the AMC-FFPE CRC was calculated using the CMSclassifier package with “SSP” method in R software^3^. All the representative pathways used for GSEA in CRC were retrieved from the study of Guinney *et al*.^3^. All the representative aberrant pathways used in HGSOC for each subtype were obtained from previous studies^48–53^.

### 3.10 Training a high-grade serous ovarian carcinoma classifier

To better classify the HGSOC cohort, we constructed a high-grade serous carcinoma classifier based on the Mayo cohort and the corresponding signatures and Mayo-defined labels^36^. Specifically, we used 147 samples (GSE53963) with 412 signatures to construct the classifier by using prediction analysis of microarray^54^. The dataset originally contained 635 probe ID signatures. Because we wanted to construct a classifier suitable for our reconstructed gene expression profile, we transferred the probes into Entrez IDs, and the Entrez IDs were then intersected with the 10,240 genes reconstructed from the network. Finally, 412 Entrez ID genes were retained. After 5-fold cross-validation, we selected threshold 0 with the minimal overall misclassification error as the final threshold setting for constructing the final classifier (**Supplementary Figure 15)**.

### 3.11 Statistical analysis

Wilcoxon tests were used for comparing groups. Detailed test settings were mentioned in the specific figures. Log-rank tests were conducted to compare overall survival for different groups in the univariate setting. Multiple testing correction was applied using the Benjamini–Hochberg method whenever required. All analyses were performed using R v 4.0.0 (www.R-project.org) and Bioconductor v 3.12. A p-value of <0.05 was considered statistically significant.

## 4. Discussion

FFPE samples often undergo severe RNA degradation, which distorts RNA-seq gene expression profiles because of reduced RNA quantity and increased fragmentation^6,55^. This long-standing challenge has limited the usability of archival FFPE transcriptomic data. To address this issue, we developed FFPERescuer, a deep learning framework for reconstructing FFPE-derived gene expression profiles and narrowing the domain gap between FF and FFPE samples. Across public and in-house datasets, our findings support three main strengths of FFPERescuer. First, the framework improved the concordance of FFPE-derived expression profiles with their reference profiles and better preserved biologically meaningful signals, indicating its ability to recover degraded transcriptomic information. Second, reconstruction enhanced the clinical relevance of downstream analyses: in colorectal and ovarian cancer, cancer subtyping, and clinical association analyses yielded results that were more concordant with biological expectations and more clinically informative. Third, FFPERescuer showed potential applicability across multiple cancer types. Although in-depth real-world analyses were performed in colorectal and ovarian cancer, simulation analyses showed consistent performance across seven cancer types, supporting its broader applicability across cancer types. Together, these findings highlight the practical value of reconstruction for unlocking the value of retrospective FFPE cohorts for cancer transcriptomics.

To our knowledge, studies that explicitly incorporate TIN, an NGS-based RNA integrity metric, into deep learning reconstruction of FFPE RNA-seq data remain limited. Because TIN can, to some extent, reflect the degree of RNA degradation, we used it to characterize degradation patterns across cancers. Notably, TIN patterns tended to be similar within the same chromosome across different cancer types, suggesting that FFPE-related RNA degradation may be selective rather than purely random. This chromosome-level consistency across cancers supports the rationale for training the autoencoder on FF TCGA pan-cancer data.

FFPE-derived RNA-seq output is influenced by multiple pre-analytical and analytical factors, including tissue handling, fixation, RNA isolation, and library preparation. However, public datasets rarely provide paired FF-FFPE RNA-seq data generated under controlled settings with defined experimental variations. Therefore, using our in-house PWH CRC paired FF-FFPE cohort, we examined the effects of two pre-analytical factors under different conditions. These analyses provide additional insight into how experimental conditions shape FFPE-derived RNA-seq profiles and complement the computational component of our study.

The PolyA-based library preparation strategy is widely used (**Supplementary Figure 11**) because of its convenience and relatively low cost, but our observations suggest that RNA-seq data generated using PolyA-based library preparation may be more vulnerable to the effects of RNA degradation. This further highlights the need for computational reconstruction of FFPE- derived RNA-seq profiles in real-world practice.

To improve reconstruction across heterogeneous datasets, we implemented a dataset-specific gene selection strategy for training the partial FF-encoder. Distinct high-TIN gene patterns across datasets (**Supplementary Figure 13**) likely reflect differences in sample handling and processing protocols. Tailoring input genes for each dataset allows better use of relatively preserved transcripts while reducing the impact of severely degraded genes.

This study has several limitations. First, although TIN is informative for RNA integrity, it does not fully capture all aspects of RNA degradation and therefore provides an imperfect reference for input-gene selection. Second, in simulation analyses, correlations between reconstructed FF samples and original FF samples did not approach 1.0, indicating that reconstruction may introduce some distortion. Future work should integrate additional degradation- aware features and further optimize model architectures to improve fidelity and generalizability.

In summary, our study provides both experimental insights and a computational solution for improving the usability of FFPE-derived RNA-seq data. Using nearly 10,000 FF TCGA samples, we developed FFPERescuer, a robust framework that reconstructs distorted FFPE transcriptomic profiles and performs effectively in both simulated and real-world datasets. By recovering biologically meaningful signals, strengthening clinically relevant downstream analyses, and showing potential applicability across multiple cancer types, FFPERescuer expands the value of archival FFPE specimens for transcriptomic research. Given the widespread availability of FFPE tissues and their rich clinical information in hospital, this framework may facilitate broader use of FFPE-derived NGS data in cancer research and translational applications.

## Abbreviations

CRC: colorectal cancer
BRCA: breast invasive carcinoma
OV: ovarian serous cystadenocarcinoma
BLCA: bladder urothelial carcinoma
LUAD: lung adenocarcinoma
GC: gastric carcinoma
MC: melanoma carcinoma
GBM: glioblastoma carcinoma
PC: prostate carcinoma
FFPE: formalin-fixed paraffin-embedded
FF: fresh frozen
OS: overall survival
rRNA: ribosomal RNA
NBF: neutral-buffered formalin
DE: differential expression
GSEA: gene set enrichment analysis
HGSOC: high-grade serous ovarian carcinoma
PolyA: poly(A)^+^ RNA
TPM: transcripts per million
TIN: transcript integrity number

## Funding

This work was funded by a grant from Shenzhen Medical Research Funds (C2303002, X.W.), a startup grant (4937084, X.W.), a direct grant (2024.175, X.W.), grants by the Faculty Postdoctoral Fellowship Scheme (FPFS/24-25/053, FPFS/23-24/061C, FPFS/23-24/060, X.W), and Research Committee – Group Research Scheme 2022-23 (WW/rc/grs2223/0560/23en, X.W.), from the Chinese University of Hong Kong, and grants from the Research Grants Council (AoE/M-401/20, R4007-23, C4024-22GF, 14104223, 11103921, and 14111522), and Health and Medical Research Fund (08192166). This work was also partially sponsored by Jiangxi Overseas High-Level Talent Project (20232BCJ25029) awarded to X.W.

## Data availability

The RNA-sequencing data of the PWH CRC cohort and AMC-FFPE CRC cohort have been deposited to the GEO database https://www.ncbi.nlm.nih.gov/geo/query/acc.cgi and assigned the identifiers GSE298630 and GSE299799, respectively. The raw sequencing data of the West China HGSOC cohort were deposited to the Genome Sequence Archive of Beijing Institute of Genomics, Chinese Academy of Sciences: HRA011750.

## Code availability

All relevant and original codes related to the article are publicly available on GitHub (https://github.com/CityUHK-CompBio/FFPE-rescuer).

## Supplementary Materials

Supplementary Figure 1 A comparison of transcript integrity number (TIN)s between paired FF and FFPE tissues.

Supplementary Figure 2 An Overview of TIN means across FF and FFPE- derived RNA-seq samples.

Supplementary Figure 3 Differences of TIN between FF CRC and FFPE CRC samples which both had total RNA library preparation protocol.

Supplementary Figure 4 An Overview of TIN means of tumor microenvironment signatures and Hallmark signatures across FF and FFPE-derived RNA-seq samples.

Supplementary Figure 5 Pearson correlations between noisy gene expression and FF gene expression in paired samples from seven cancer types.

Supplementary Figure 6 An Overview of TIN means across four focused FFPE-derived RNA-seq datasets.

Supplementary Figure 7 An Overview of TINs of cancer-specific microenvironment signatures.

Supplementary Figure 8 A comparison of gene expression levels of paired FF samples and real-world CRC FFPE samples before and after applying FFPERescuer.

Supplementary Figure 9 A comparison of gene expression level of real- world CRC FFPE samples before and after applying FFPERescuer.

Supplementary Figure 10 Sample size distribution of FFPE tumor samples among nine cancer types.

Supplementary Figure 11 Comparison of sample size of excluded samples with distinct library preparation methods.

Supplementary Figure 12 Relationship between gene expression (TPM) and TIN means in FFPE-derived RNA-seq samples.

Supplementary Figure 13 Explanations for the strategies used in constructing the FFPERescuer.

Supplementary Figure 14 Explanations for the type of noise added in the simulation analysis.

Supplementary Figure 15 Evaluation on the trained HGSOC classifier.

Supplementary Table 1 Specimen information of the PWH CRC cohort.

Supplementary Table 2 Significant pathways for genes with low TINs using GSEA.

Supplementary Table 3 Significant pathways for genes with high TINs using GSEA.

Supplementary Table 4 CMS of the 12 FF-FFPE-matched samples (AMC- FFPE CRC).

Supplementary Table 5 Subtype of the 27 FFPE HGSOC samples (West China HGSOC).

Supplementary Table 6 A summary of public FFPE datasets.

Supplementary Table 7 A summary of the validation datasets.

Supplementary Table 8 A summary of public FF datasets used.

Supplementary Table 9 FF TCGA samples used for GSEA in simulation experiments.

## Notes

### Competing Interest Statement

The authors have declared no competing interest.

