## Supplementary figures for "FFPERescuer: deep unsupervised domain adaptation for the reconstruction of gene expression profiles derived from formalin-fixed paraffin-embedded samples"

[1]Department of Surgery, The Chinese University of Hong Kong, Shatin, Hong Kong SAR, China.
[2]Institute of Digestive Disease, The Chinese University of Hong Kong, Shatin, Hong Kong SAR, China.
[3]Department of Obstetrics and Gynecology, Key Laboratory of Birth Defects and Related Diseases of Women and Children of MOE, State Key Laboratory of Biotherapy, West China Second Hospital, Sichuan University and Collaborative Innovation Center, Chengdu, China.
[4]Laboratory for Experimental Oncology and Radiobiology, Center for Experimental and Molecular Medicine, Cancer Center Amsterdam, Amsterdam Gastroenterology Endocrinology Metabolism, Amsterdam University Medical Centers, Amsterdam, The Netherlands.
[5]Oncode Institute, Amsterdam, The Netherlands.
[6]Department of General Surgery (Department of Colorectal Surgery), The Sixth Affiliated Hospital, Sun Yat-sen University, Guangzhou, China.
[7]Shenzhen Research Institute, The Chinese University of Hong Kong, Shenzhen, China.
[8]Li Ka Shing Institute of Health Sciences, The Chinese University of Hong Kong, Shatin, Hong Kong SAR, China.
**[*]Correspondence**

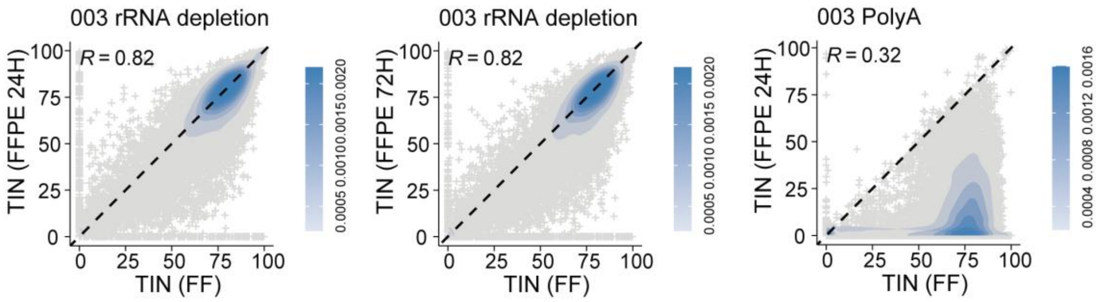


**Supplementary Figure 1 A comparison of transcript integrity number (TIN)s between paired FF and FFPE tissues.**

Scatter plots showing the Pearson correlation between whole-genome TINs from FF tissue and FFPE tissue for the same patient.

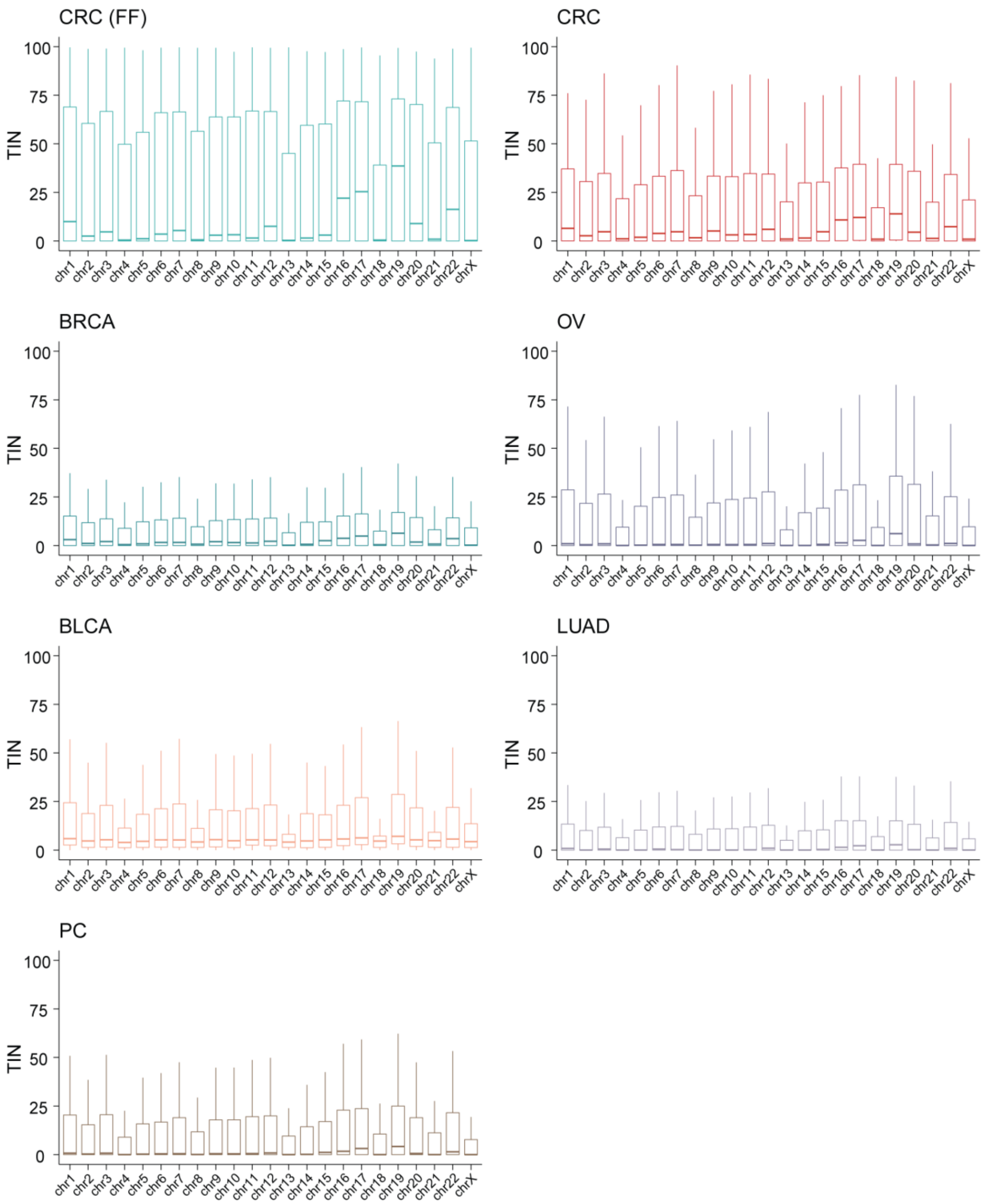


**Supplementary Figure 2 An Overview of TIN means across FF and FFPE-derived RNA-seq samples.**

Boxplots showing the TIN means of seven types of cancer from FFPE RNA-seq samples and one type of cancer from FF RNA-seq samples, categorized by chromosomal location.

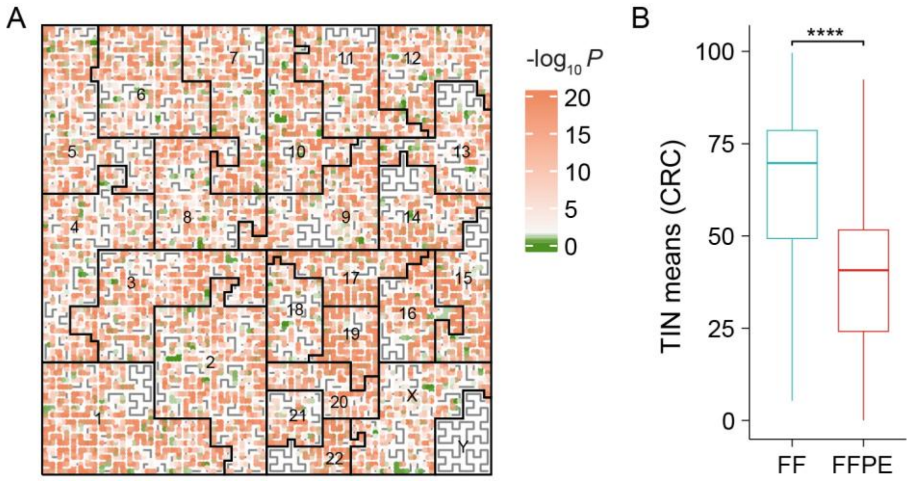


**Supplementary Figure 3 Differences of TIN between FF CRC and FFPE CRC samples which both had total RNA library preparation protocol.**

**A**, Hilbert curve illustrating the significant values of whole-genome TIN mean difference test between 145 CRC FF transcriptomes and 75 CRC FFPE transcriptomes. **B,** Boxplot showing the whole-genome TIN means of the CRC samples used in **Supplementary Figure 3A**. Adjusted $P \leq 0.0001$ (****). The center line of the boxplots indicates the median, the edges indicate the interquartile range. In this figure, all the p-values were derived from unpaired one-sided Student’s t-tests and were Benjamini–Hochberg adjusted.

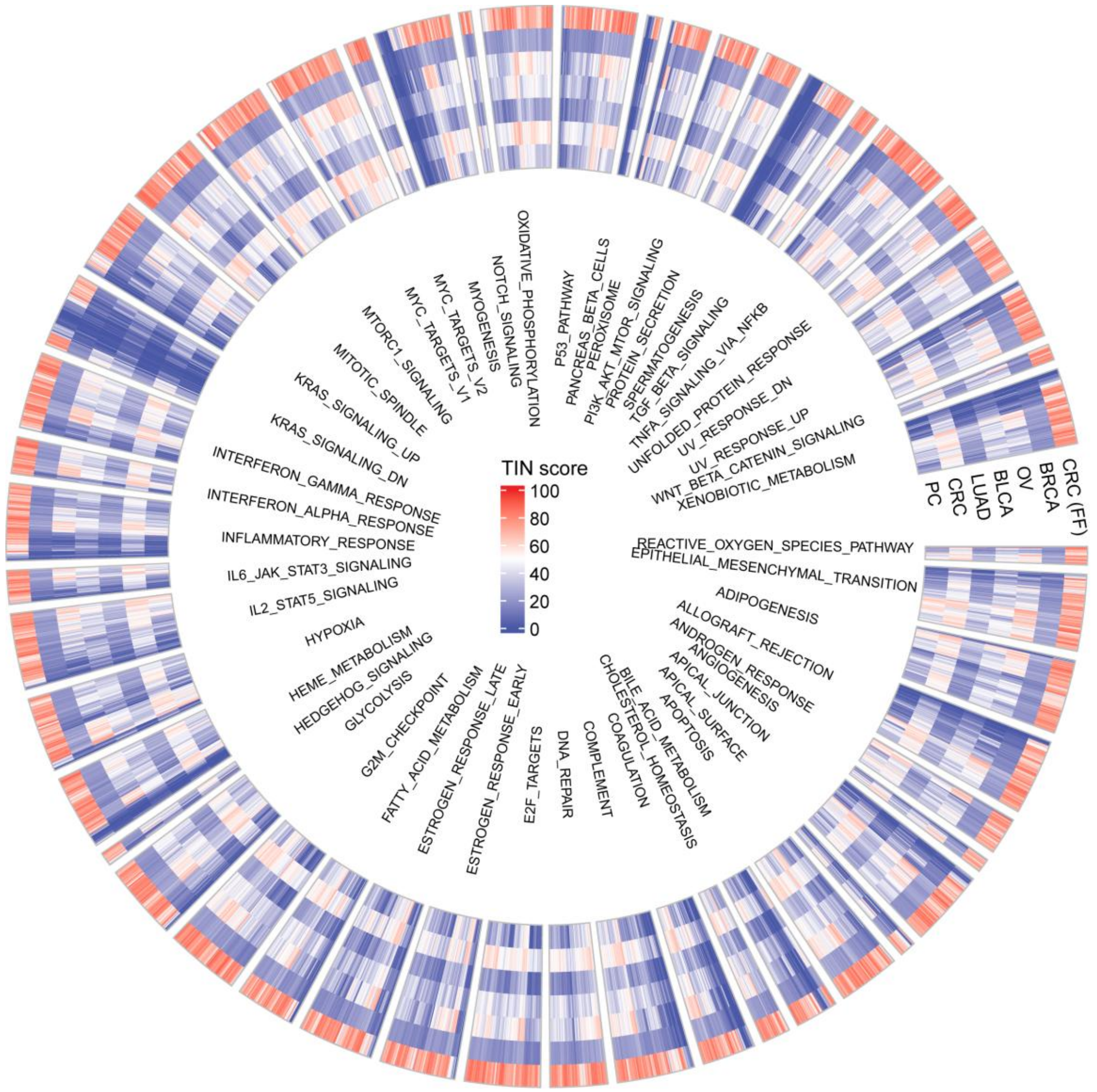


**Supplementary Figure 4 An Overview of TIN means of tumor microenvironment signatures and Hallmark signatures across FF and FFPE-derived RNA-seq samples.** Circos heatmap illustrating the TIN means of the genes from the 50 Hallmark gene sets.

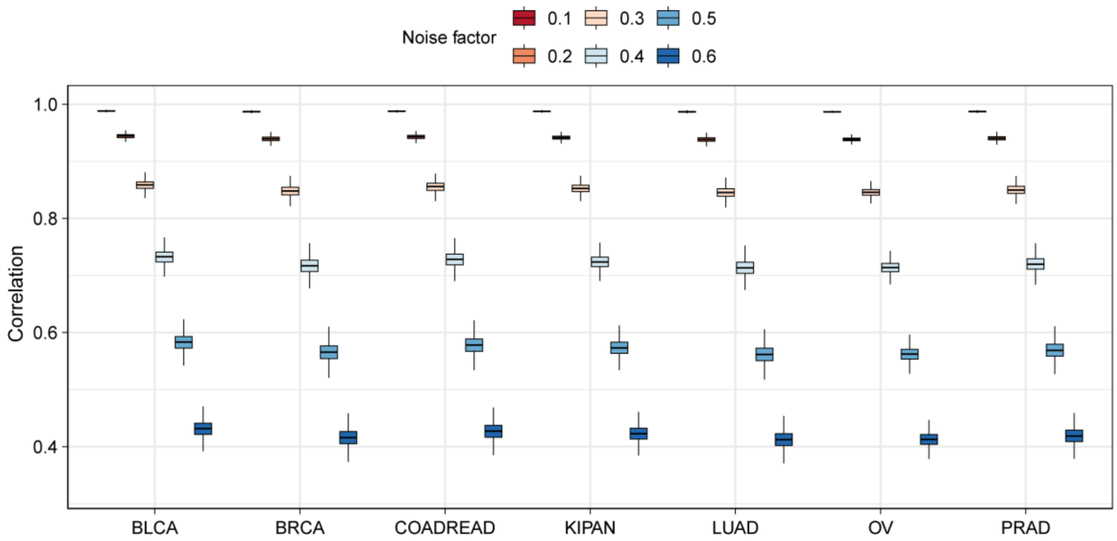


**Supplementary Figure 5 Pearson correlations between noisy gene expression and FF gene expression in paired samples from seven cancer types.**

Boxplot showing the Pearson correlations between noisy gene expression and FF gene expression in paired samples from seven cancer types. Six sets of noisy gene expression profiles were generated with increasing noise (noise factor equals to 0.1, 0.2, 0.3, 0.4, 0.5, and 0.6).

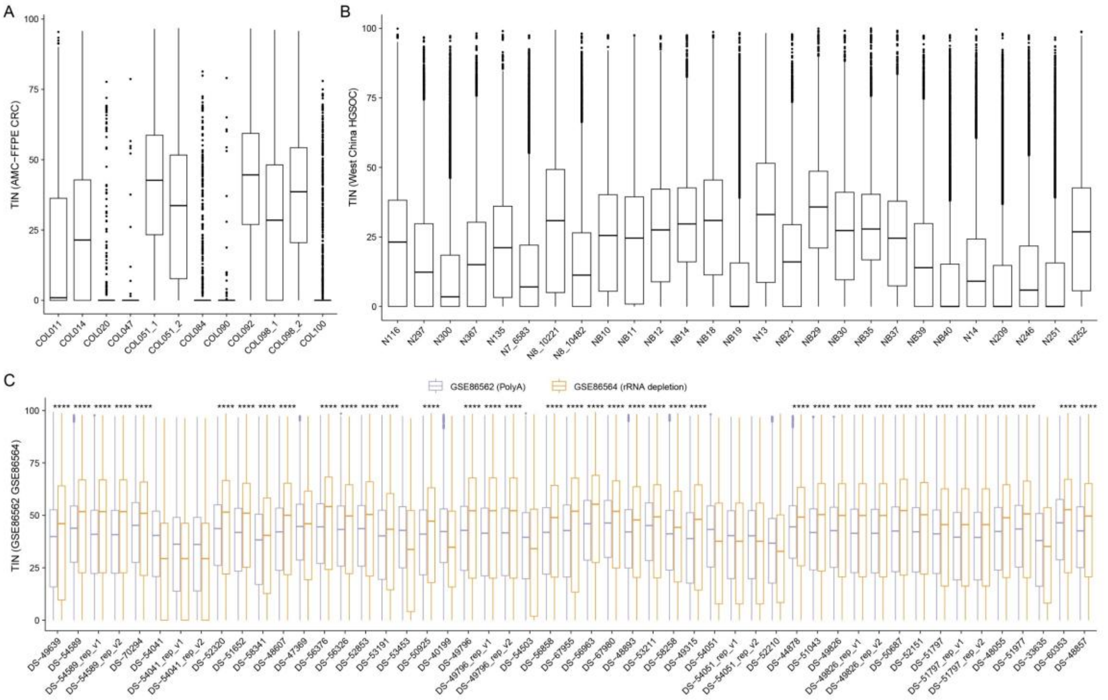


**Supplementary Figure 6 An Overview of TIN means across four focused FFPE-derived RNA-seq datasets.**

**A–C,** Boxplots showing the whole-genome TIN means of the AMC-FFPE CRC (**A**), the West China HGSOC (**B**). The sample sizes are as follows: AMC-FFPE CRC, N = 12; West China HGSOC, N = 27. **C,** Boxplots showing the whole-genome TIN means of paired FFPE samples with different library preparation methods (**Supplementary Table 4**). The sample sizes are both 51. The statistical differences were derived from paired one-sided Student's t-tests and adjusted by Benjamini–Hochberg method; Adjusted $P \leq 0.0001$ (****); Adjusted $P \leq 0.05$ (*).

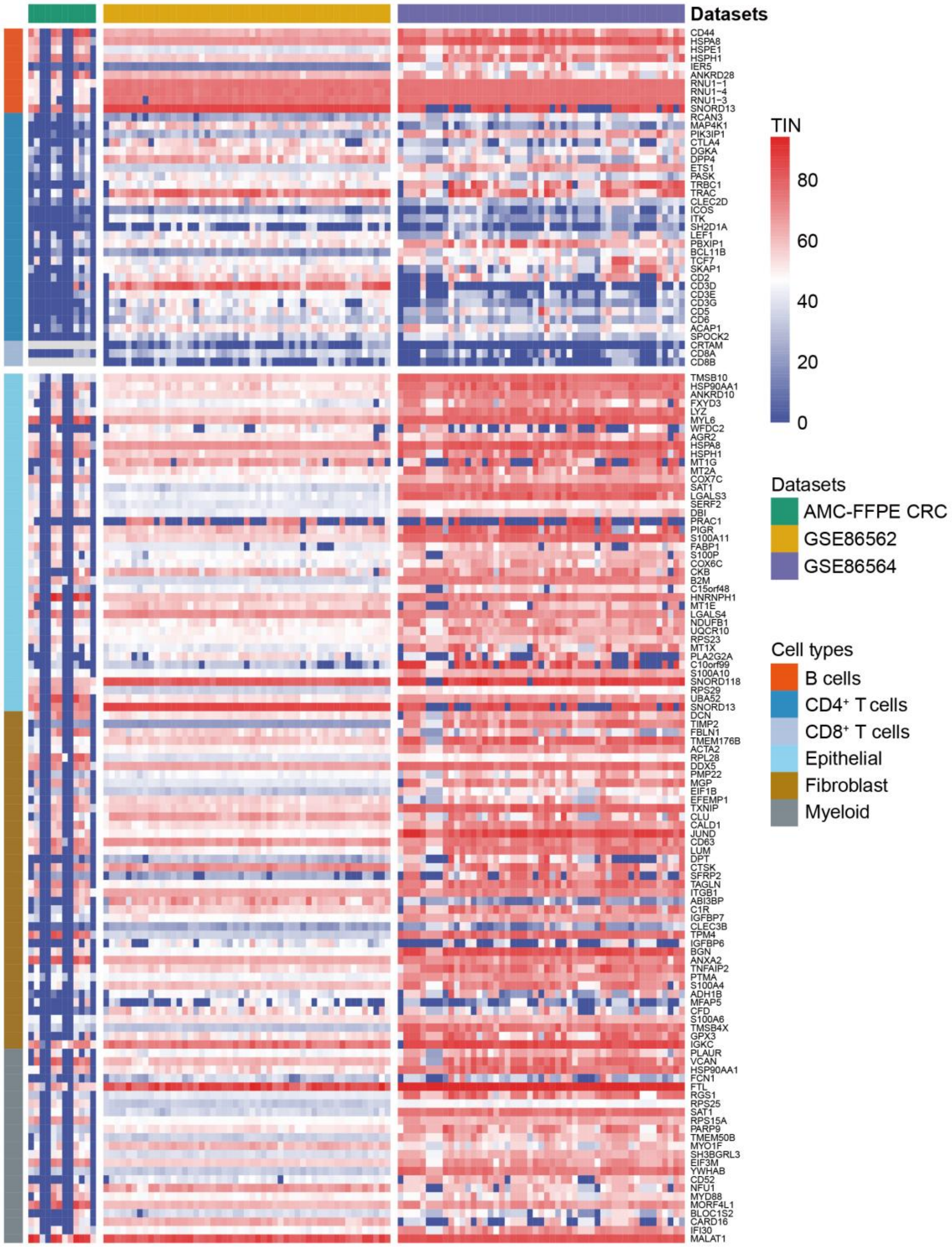


**Supplementary Figure 7 An Overview of TINs of cancer-specific microenvironment signatures.**

Heatmap showing the TINs of CRC signature genes in the AMC-FFPE CRC, GSE86562, and GSE86564.

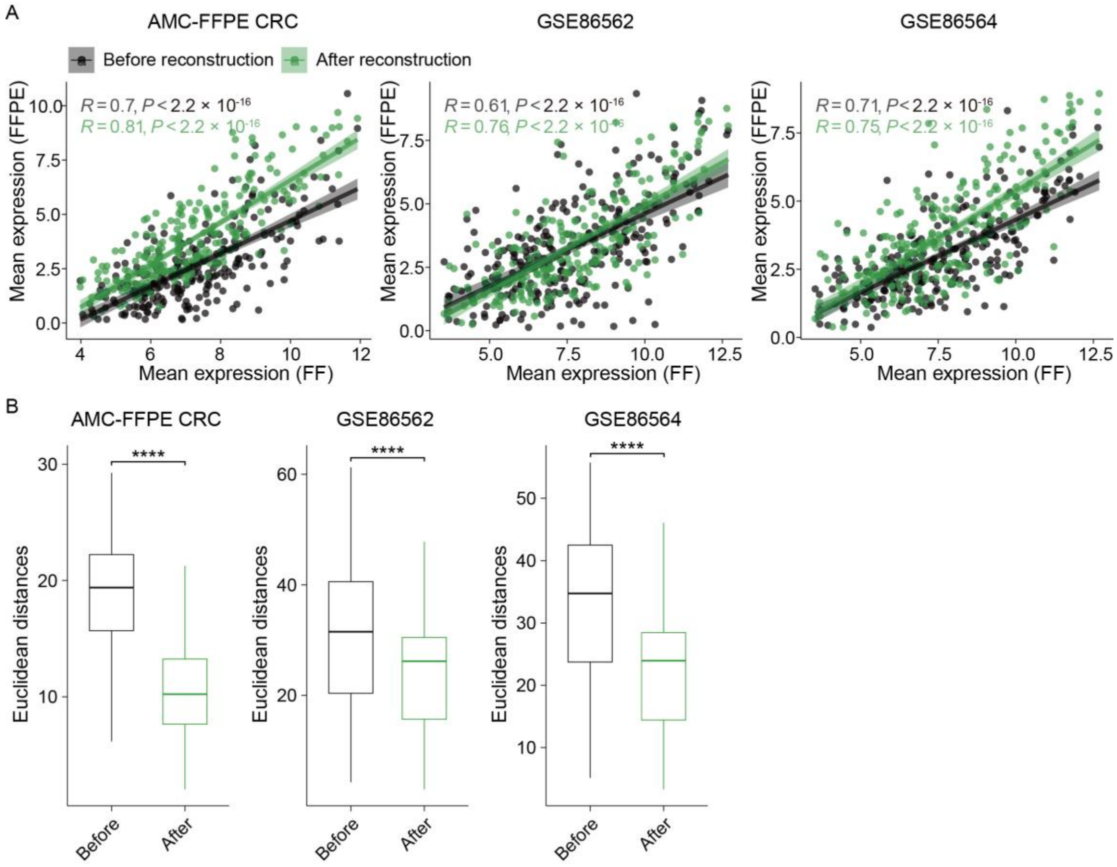


**Supplementary Figure 8 A comparison of gene expression levels of paired FF samples and real-world CRC FFPE samples before and after applying FFPERescuer.** **A,** Scatterplots showing 273 CRC gene signature expression of paired FF and FFPE samples from the AMC-FFPE CRC (left), GSE86562 (middle), and GSE86564 (right) before and after reconstruction. **B**, Boxplot showing the pair-wise non-immune cell signatures (Myeloid cell, epithelial cell, and fibroblast cell) Euclidean distances of FF gene expression and FFPE gene expression among three datasets before and after reconstruction. The statistical differences were derived from paired one-sided Wilcoxon signed-rank tests; Adjusted $P \leq 0.0001$ (****). The center line of the boxplots indicates the median, the edges indicate the interquartile range. All the p-values were Benjamini–Hochberg adjusted.

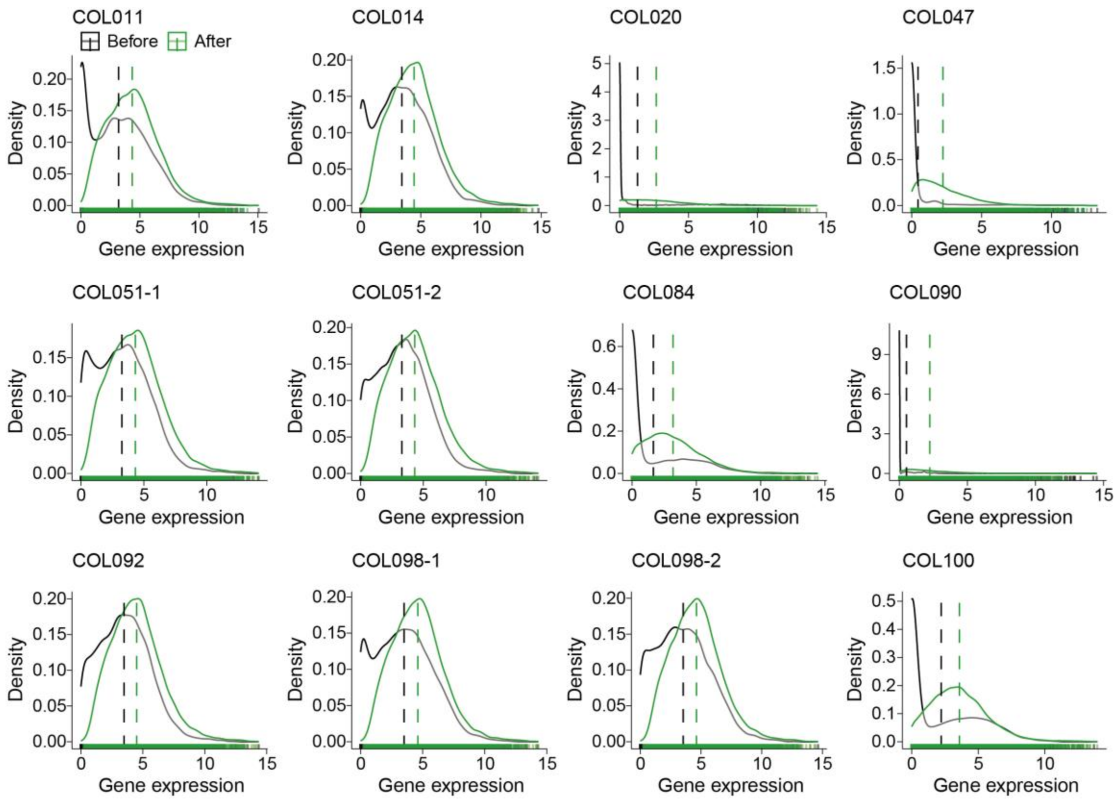


**Supplementary Figure 9 A comparison of gene expression level of real-world CRC FFPE samples before and after applying FFPERescuer.**

Density plots showing the whole-genome gene expression distribution of each FFPE sample before and after reconstruction on the AMC-FFPE CRC (N = 12).

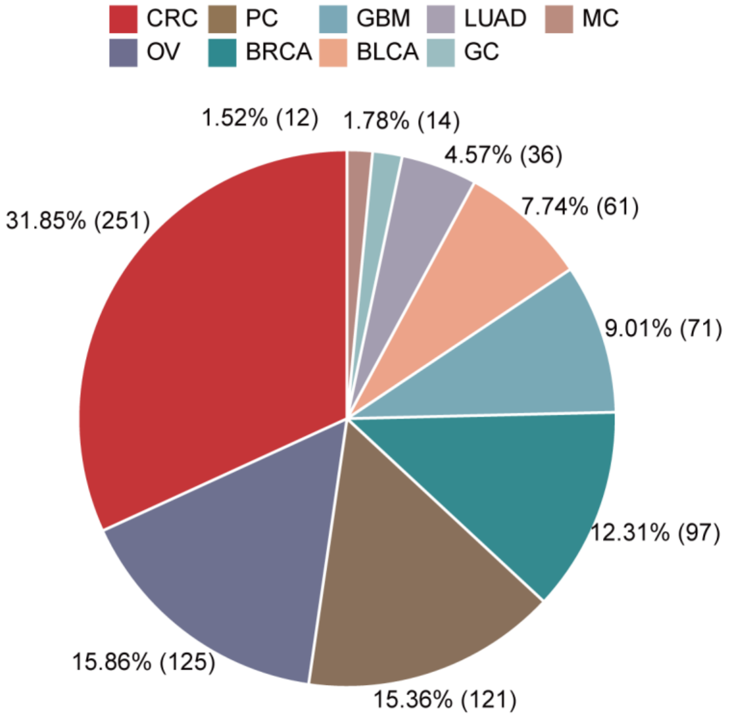


**Supplementary Figure 10 Sample size distribution of FFPE tumor samples among nine cancer types.**

Pie chart illustrating all the sample size and proportion of each type of cancer from FFPE RNA-seq the study has collected (N = 788). The number of samples in each type was labeled.

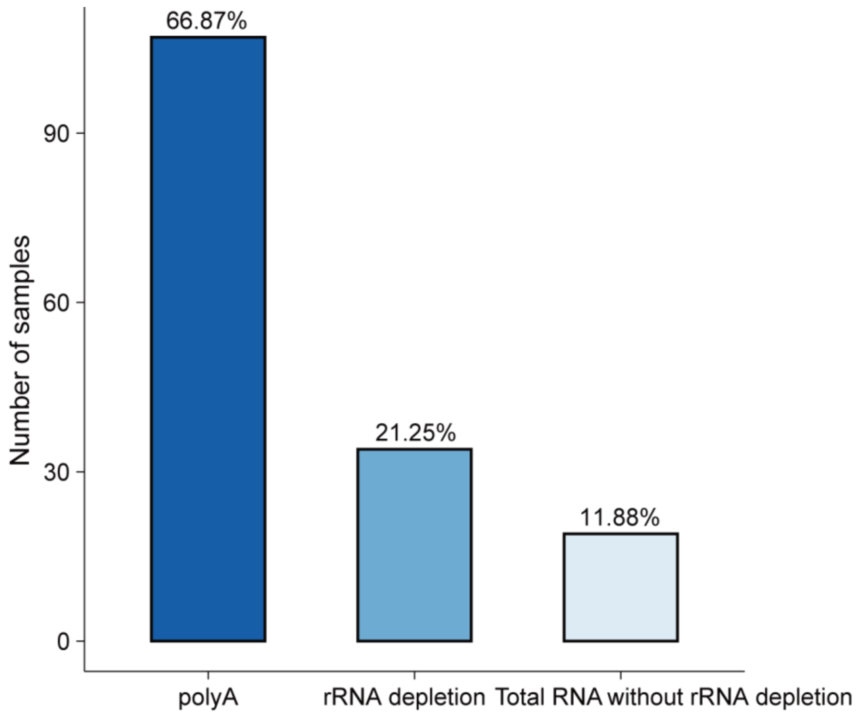


**Supplementary Figure 11 Comparison of sample size of excluded samples with distinct library preparation methods.**

Bar plot showing the number of samples filtered out due to many zero TIN mean genes in one sample. Proportion of each type was labeled. Total RNA means the rRNA was not removed.

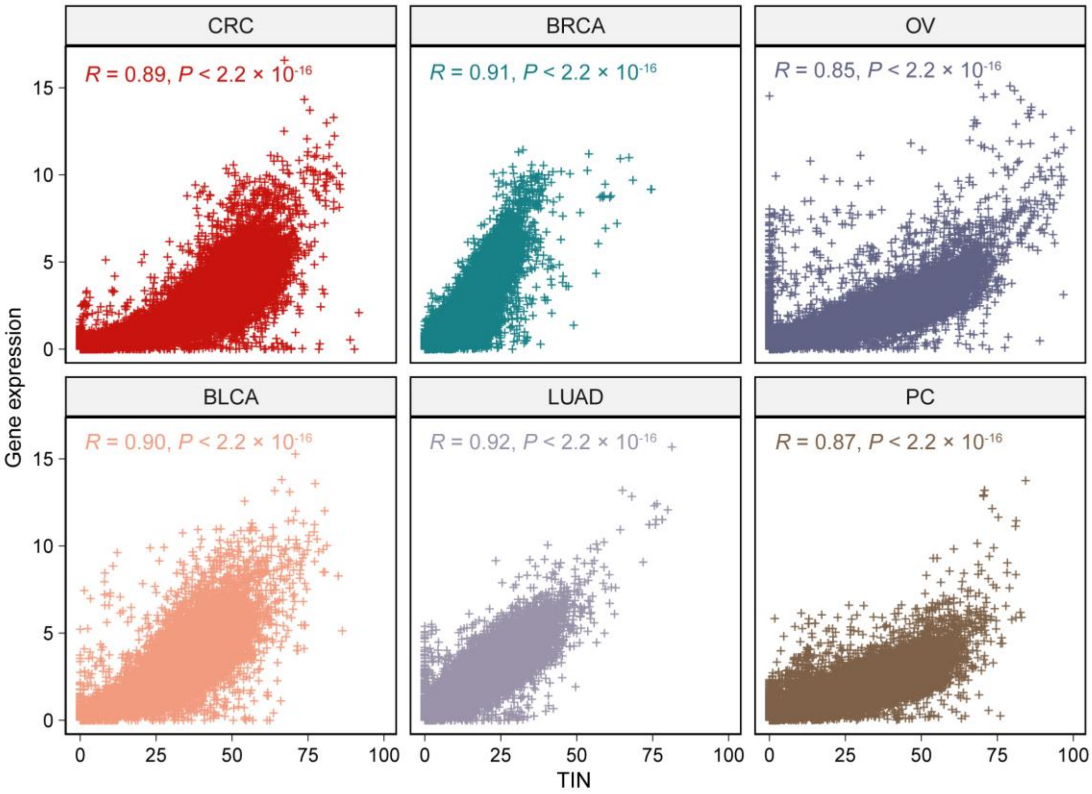


**Supplementary Figure 12 Relationship between gene expression (TPM) and TIN means in FFPE-derived RNA-seq samples.**

Scatterplots showing the Pearson correlation of the paired gene expression means and TIN means of each gene within each cancer type. R represents the Pearson correlation coefficient. The p-values were Benjamini–Hochberg adjusted.

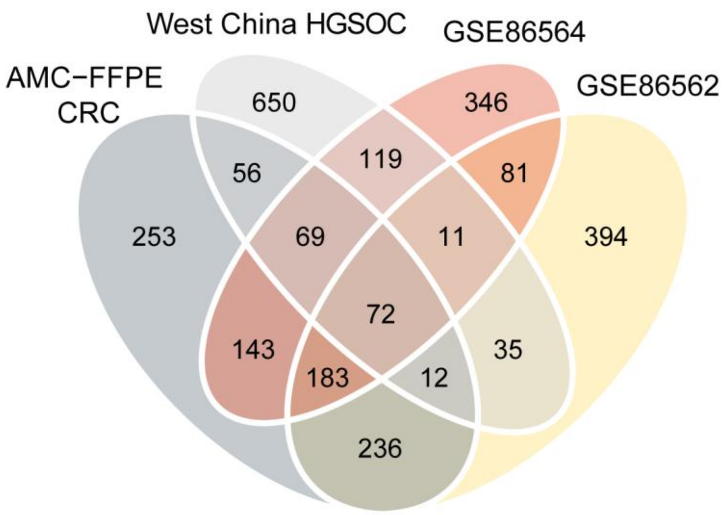


**Supplementary Figure 13 Explanations for the strategies used in constructing the FFPERescuer.**

Venn diagram showing overlaps for cohorts with the same genes whose TIN median are bigger than 20.

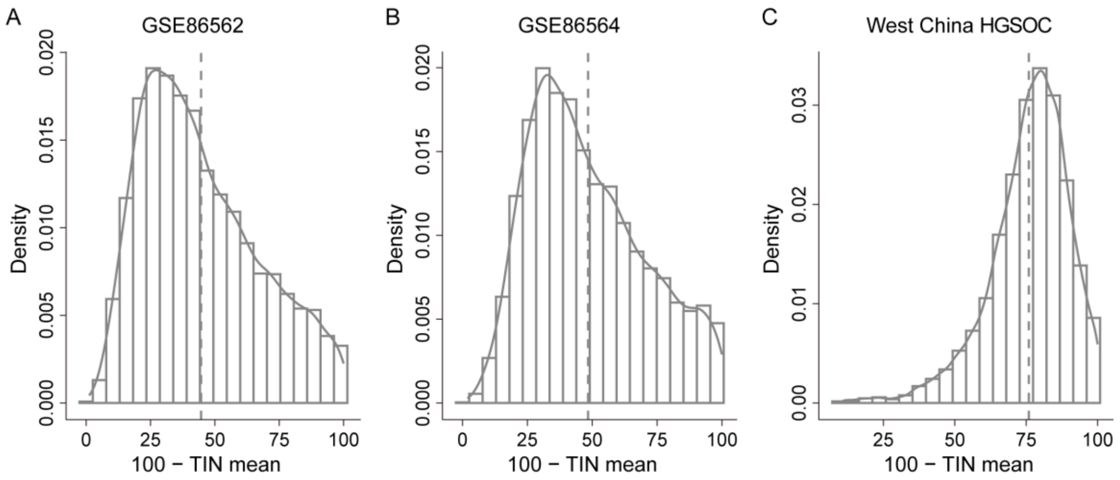


**Supplementary Figure 14 Explanations for the type of noise added in the simulation analysis.**

**A–C**, Density plot illustrating the estimated degradation (100 – TIN mean) of three representative datasets: GSE86562 (**A**), GSE86562 (**B**), and the West China HGSOC (**C**) across the whole-genome.

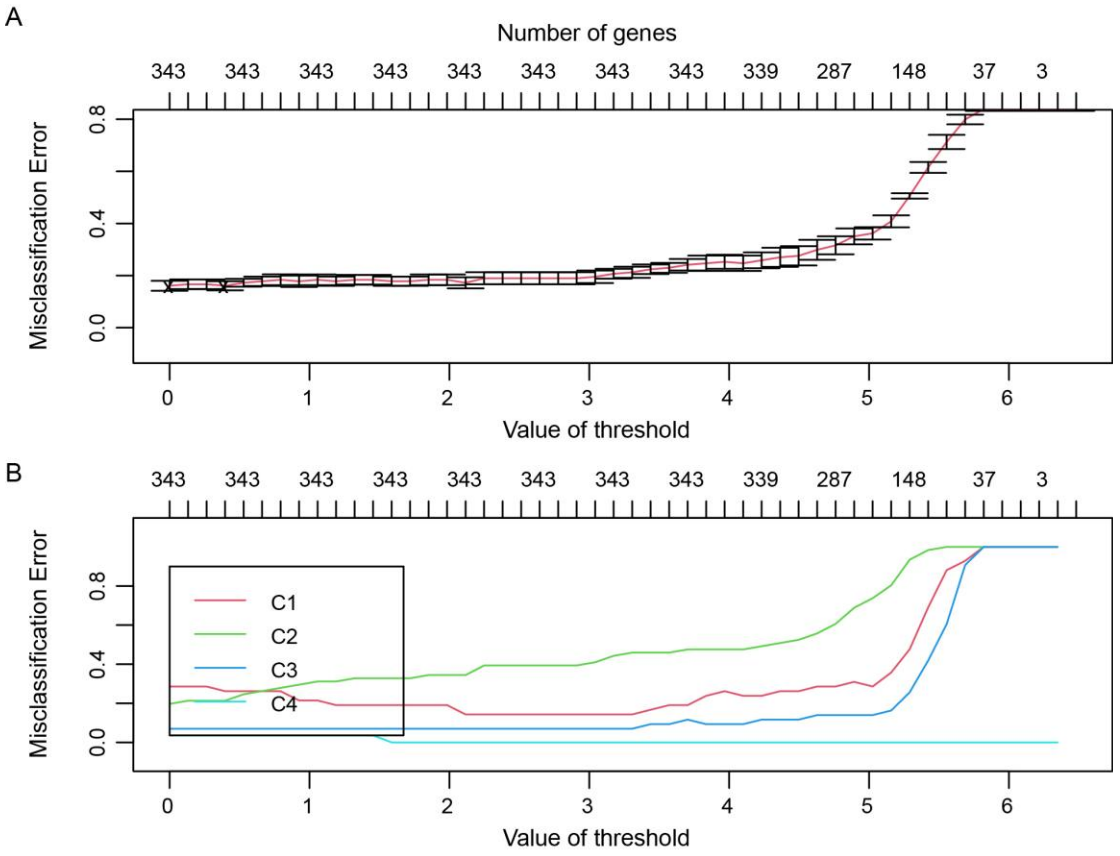


**Supplementary Figure 15 Evaluation on the trained HGSOC classifier.**

**A**–**B,** Averaged 5-fold cross-validated error curves (**A**) and class-wise 5-fold cross-validated error curves (**B**) based on shrunken centroid classifier with different thresholds and different number of genes. The training procedure was conducted with the pamr R package.

**Supplementary Table 1 Specimen information of the PWH CRC cohort.**

| ID* | Status$ | RNA (ng/uL) | A260/A280 | Volume | Ribosomal RNA depletion + RNA-sequencing | PolyA enrichment + RNA-sequencing | Platform |
|---|---|---|---|---|---|---|---|
| FF_003 | RNA ready, -80C | 260.1 | 2.08 | 70uL | 1 | 1 | Illumina NovaSeq 6000 |
| FFPE_003_24_h | RNA ready, -80C | 341.7 | 2.08 | 30uL | 1 | 1 | Illumina NovaSeq 6000 |
| FFPE_003_72_h | RNA ready, -80C | 237 | 2.03 | 30uL | 1 | | Illumina NovaSeq 6000 |

**Supplementary Table 2 Significant pathways for genes with low TINs using GSEA.**

| ID | Set size | Enrichment score | NES | P-value | Adjusted *P* | Q-value | Rank | Leading edge |
|---|---|---|---|---|---|---|---|---|
| WEBER_METHYLATED_ICP_IN_SPERM_DN | 11 | -0.582444438 | -1.97123316 | 0.003623188 | 0.00454894 | 0.000445229 | 19201 | tags=82%, list=38%, signal=51% |
| GOBP_NEGATIVE_REGULATION_OF_VASCULAR_ENDOTHELIAL_GROWTH_FACTOR_PRODUCTION | 9 | -0.812840367 | -2.55449814 | 0.00330033 | 0.005508483 | 0.000942046 | 9461 | tags=100%, list=19%, signal=81% |
| GOMF_TYPE_I_INTERFERON_RECEPTOR_BINDING | 17 | -0.554593492 | -2.24309687 | 0.005 | 0.007817521 | 0.001336932 | 19677 | tags=88%, list=39%, signal=54% |
| HP_Y_LINKED_INHERITANCE | 17 | -0.546348322 | -2.20974864 | 0.005 | 0.007817521 | 0.001336932 | 15250 | tags=76%, list=30%, signal=53% |
| GOMF_GLUTAMATE_RECEPTOR_ACTIVITY | 26 | -0.374076908 | -1.84725653 | 0.006289308 | 0.009335577 | 0.001596545 | 29128 | tags=92%, list=58%, signal=39% |
| REACTOME_BETA_DEFENSINS | 35 | -0.534794915 | -2.86830996 | 0.008333333 | 0.009900963 | 0.000969059 | 13869 | tags=74%, list=27%, signal=54% |
| GOMF_NEUROPEPTIDE_ACTIVITY | 30 | -0.410153996 | -2.0788509 | 0.007042254 | 0.010279429 | 0.00175796 | 29286 | tags=97%, list=58%, signal=41% |

| | | | | | | | | |
|---|---|---|---|---|---|---|---|---|
| REACTOME_DEFENSINS | 45 | -0.421555936 | -2.53865182 | 0.010204082 | 0.012026892 | 0.001177135 | 13869 | tags=62%, list=27%, signal=45% |
| GOMF_NEUROPEPTIDE_RECEPTOR_ACTIVITY | 47 | -0.347489895 | -2.10931407 | 0.009433962 | 0.013459343 | 0.002301781 | 29924 | tags=89%, list=59%, signal=36% |
| GOMF_NEUROTRANSMITTER_RECEPTOR_ACTIVITY_INVOLVED_IN_REGULATION_OF_POSTSYNAPTIC_MEMBRANE_POTENTIAL | 55 | -0.30337083 | -2.05991648 | 0.012345679 | 0.016860803 | 0.002883489 | 29517 | tags=84%, list=58%, signal=35% |
| GOMF_POSTSYNAPTIC_NEUROTRANSMITTER_RECEPTOR_ACTIVITY | 70 | -0.296870936 | -2.13637622 | 0.018181818 | 0.023541286 | 0.004025968 | 29629 | tags=81%, list=59%, signal=34% |
| GOMF_EXTRACELLULAR_LIGAND_GATED_MONOATOMIC_ION_CHANNEL_ACTIVITY | 73 | -0.234800824 | -1.71188527 | 0.018867925 | 0.02436562 | 0.004166943 | 29128 | tags=78%, list=58%, signal=33% |
| WEBER_METHYLATED_ICP_IN_FIBROBLAST | 19 | -0.439544495 | -1.96251541 | 0.021929825 | 0.02495332 | 0.002442312 | 19201 | tags=89%, list=38%, signal=55% |
| GOMF_CCR6_CHEMOKINE_RECEPTOR_BINDING | 7 | -0.631426343 | -1.75205715 | 0.020710059 | 0.026449388 | 0.004523303 | 15604 | tags=71%, list=31%, signal=49% |
| GOMF_TRACE_AMINE_RECEPTOR_ACTIVITY | 6 | -0.645108911 | -1.62692655 | 0.021021021 | 0.026750366 | 0.004574776 | 17929 | tags=100%, list=35%, signal=65% |

| | | | | | | | | |
|---|---|---|---|---|---|---|---|---|
| GOCC_KERATIN_FILAMENT | 90 | -0.226336947 | -1.71479798 | 0.033333333 | 0.041126905 | 0.0070334 13 | 22323 | tags=71%, list=44%, signal=40% |
| MIKKELSEN_ES_LCP_WITH_H3 K27ME3 | 13 | -0.471336433 | -1.7071709 | 0.037593985 | 0.041810889 | 0.0040922 51 | 22950 | tags=77%, list=45%, signal=42% |
| GOMF_NEUROTRANSMITTER_ RECEPTOR_ACTIVITY | 97 | -0.234778978 | -1.78862859 | 0.04 | 0.048556218 | 0.0083039 54 | 29629 | tags=79%, list=59%, signal=33% |

**Supplementary Table 3 Significant pathways for genes with high TINs using GSEA.**

| ID | Set size | Enrichment score | NES | P-value | Adjusted *P* | Q-value | Rank | Leading edge |
|---|---|---|---|---|---|---|---|---|
| HALLMARK_MYC_TARGETS_V1 | 195 | 0.808581441 | 3.7254 | 0.001 | 0.00124794 | 5.25449E-05 | 6062 | tags=87%, list=12%, signal=77% |
| HALLMARK_OXIDATIVE_PHOSPHORYLATION | 185 | 0.770220405 | 3.5351 | 0.001001 | 0.00124794 | 5.25449E-05 | 9987 | tags=92%, list=20%, signal=74% |
| HALLMARK_UNFOLDED_PROTEIN_RESPONSE | 108 | 0.786887029 | 3.47238 | 0.001017 | 0.00124794 | 5.25449E-05 | 8346 | tags=90%, list=17%, signal=75% |
| HALLMARK_MTORC1_SIGNALING | 195 | 0.743958282 | 3.42766 | 0.001 | 0.00124794 | 5.25449E-05 | 10712 | tags=87%, list=21%, signal=69% |
| HALLMARK_EPITHELIAL_MESENCHYMAL_TRANSITION | 197 | 0.736516312 | 3.39524 | 0.001001 | 0.00124794 | 5.25449E-05 | 9901 | tags=77%, list=20%, signal=62% |
| HALLMARK_MITOTIC_SPINDLE | 198 | 0.736057176 | 3.39518 | 0.001001 | 0.00124794 | 5.25449E-05 | 11217 | tags=88%, list=22%, signal=69% |
| HALLMARK_ANDROGEN_RESPONSE | 97 | 0.764900825 | 3.33785 | 0.00102 | 0.00124794 | 5.25449E-05 | 10067 | tags=89%, list=20%, signal=71% |
| HALLMARK_P53_PATHWAY | 194 | 0.723557197 | 3.32946 | 0.001 | 0.00124794 | 5.25449E-05 | 12035 | tags=85%, list=24%, signal=65% |
| HALLMARK_DNA_REPAIR | 148 | 0.737521132 | 3.32711 | 0.001005 | 0.00124794 | 5.25449E-05 | 10489 | tags=85%, list=21%, signal=68% |
| HALLMARK_APOPTOSIS | 160 | 0.731488845 | 3.31509 | 0.001002 | 0.00124794 | 5.25449E-05 | 9818 | tags=77%, list=19%, signal=62% |
| HALLMARK_INTERFERON_GAMMA_RESPONSE | 196 | 0.718855597 | 3.31356 | 0.001 | 0.00124794 | 5.25449E-05 | 11522 | tags=80%, list=23%, signal=62% |
| HALLMARK_TNFA_SIGNALING_VIA_NFKB | 198 | 0.717128818 | 3.30787 | 0.001001 | 0.00124794 | 5.25449E-05 | 11196 | tags=79%, list=22%, signal=62% |

| | | | | | | | | |
|---|---|---|---|---|---|---|---|---|
| HALLMARK_PROTEIN_SECRETION | 95 | 0.761151787 | 3.30624 | 0.001027 | 0.00124794 | 5.25449E-05 | 9946 | tags=89%, list=20%, signal=72% |
| HALLMARK_UV_RESPONSE_DN | 141 | 0.733834392 | 3.30187 | 0.001004 | 0.00124794 | 5.25449E-05 | 11473 | tags=86%, list=23%, signal=67% |
| HALLMARK_ADIPOGENESIS | 192 | 0.716108558 | 3.29219 | 0.001 | 0.00124794 | 5.25449E-05 | 11542 | tags=83%, list=23%, signal=65% |
| HALLMARK_INTERFERON_ALPHA_RESPONSE | 95 | 0.756054681 | 3.2841 | 0.001027 | 0.00124794 | 5.25449E-05 | 10591 | tags=86%, list=21%, signal=68% |
| HALLMARK_G2M_CHECKPOINT | 192 | 0.713298163 | 3.27927 | 0.001 | 0.00124794 | 5.25449E-05 | 12519 | tags=86%, list=25%, signal=65% |
| HALLMARK_HYPOXIA | 194 | 0.70130772 | 3.22708 | 0.001 | 0.00124794 | 5.25449E-05 | 10657 | tags=72%, list=21%, signal=57% |
| HALLMARK_E2F_TARGETS | 195 | 0.696237222 | 3.20779 | 0.001 | 0.00124794 | 5.25449E-05 | 14690 | tags=93%, list=29%, signal=66% |
| HALLMARK_GLYCOLYSIS | 198 | 0.69517285 | 3.2066 | 0.001001 | 0.00124794 | 5.25449E-05 | 11131 | tags=75%, list=22%, signal=59% |
| HALLMARK_TGF_BETA_SIGNALING | 54 | 0.786567311 | 3.17442 | 0.001095 | 0.00124794 | 5.25449E-05 | 6924 | tags=83%, list=14%, signal=72% |
| HALLMARK_ESTROGEN_RESPONSE_EARLY | 194 | 0.688406348 | 3.16771 | 0.001 | 0.00124794 | 5.25449E-05 | 13575 | tags=78%, list=27%, signal=58% |
| HALLMARK_UV_RESPONSE_UP | 155 | 0.701231237 | 3.16669 | 0.001006 | 0.00124794 | 5.25449E-05 | 9987 | tags=68%, list=20%, signal=55% |
| HALLMARK_FATTY_ACID_METABOLISM | 156 | 0.695584344 | 3.14063 | 0.001005 | 0.00124794 | 5.25449E-05 | 10435 | tags=74%, list=21%, signal=59% |
| HALLMARK_MYC_TARGETS_V2 | 57 | 0.76407305 | 3.11587 | 0.001087 | 0.00124794 | 5.25449E-05 | 11354 | tags=96%, list=22%, signal=75% |

| | | | | | | | | |
|---|---|---|---|---|---|---|---|---|
| HALLMARK_APICAL_JUNCTION | 198 | 0.675165804 | 3.11431 | 0.001001 | 0.00124794 | 5.25449E-05 | 8094 | tags=59%, list=16%, signal=49% |
| HALLMARK_PI3K_AKT_MTOR_SIGNALING | 104 | 0.704124797 | 3.09692 | 0.001019 | 0.00124794 | 5.25449E-05 | 12869 | tags=86%, list=25%, signal=64% |
| HALLMARK_PEROXISOME | 104 | 0.702937625 | 3.0917 | 0.001019 | 0.00124794 | 5.25449E-05 | 10017 | tags=72%, list=20%, signal=58% |
| HALLMARK_IL2_STAT5_SIGNALING | 196 | 0.670392854 | 3.09017 | 0.001 | 0.00124794 | 5.25449E-05 | 12854 | tags=74%, list=25%, signal=55% |
| HALLMARK_COMPLEMENT | 200 | 0.66671604 | 3.07805 | 0.001 | 0.00124794 | 5.25449E-05 | 10171 | tags=63%, list=20%, signal=51% |
| HALLMARK_ESTROGEN_RESPONSE_LATE | 194 | 0.660050758 | 3.03723 | 0.001 | 0.00124794 | 5.25449E-05 | 13883 | tags=75%, list=27%, signal=54% |
| HALLMARK_CHOLESTEROL_HOMEOSTASIS | 73 | 0.723476301 | 3.03332 | 0.00106 | 0.00124794 | 5.25449E-05 | 12815 | tags=90%, list=25%, signal=68% |
| HALLMARK_HEME_METABOLISM | 194 | 0.652426509 | 3.00215 | 0.001 | 0.00124794 | 5.25449E-05 | 10168 | tags=65%, list=20%, signal=52% |
| HALLMARK_KRAS_SIGNALING_UP | 198 | 0.64745306 | 2.98648 | 0.001001 | 0.00124794 | 5.25449E-05 | 12668 | tags=68%, list=25%, signal=51% |
| HALLMARK_XENOBIOTIC_METABOLISM | 197 | 0.628902071 | 2.89915 | 0.001001 | 0.00124794 | 5.25449E-05 | 12490 | tags=63%, list=25%, signal=48% |
| HALLMARK_REACTIVE_OXYGEN_SPECIES_PATHWAY | 47 | 0.728775207 | 2.86648 | 0.00112 | 0.00124794 | 5.25449E-05 | 12193 | tags=87%, list=24%, signal=66% |
| HALLMARK_COAGULATION | 138 | 0.635944575 | 2.84953 | 0.001009 | 0.00124794 | 5.25449E-05 | 9566 | tags=55%, list=19%, signal=45% |
| HALLMARK_ALLOGRAFT_REJECTION | 196 | 0.613090532 | 2.82604 | 0.001 | 0.00124794 | 5.25449E-05 | 13874 | tags=65%, list=27%, signal=47% |

| | | | | | | | | |
|---|---|---|---|---|---|---|---|---|
| HALLMARK_IL6_JAK_STAT3_SIGNALING | 87 | 0.654852785 | 2.81953 | 0.001032 | 0.00124794 | 5.25449E-05 | 12744 | tags=69%, list=25%, signal=52% |
| HALLMARK_INFLAMMATORY_RESPONSE | 199 | 0.605506241 | 2.79342 | 0.001 | 0.00124794 | 5.25449E-05 | 15356 | tags=71%, list=30%, signal=50% |
| HALLMARK_MYOGENESIS | 197 | 0.604181749 | 2.7852 | 0.001001 | 0.00124794 | 5.25449E-05 | 11337 | tags=54%, list=22%, signal=42% |
| HALLMARK_WNT_BETA_CATENIN_SIGNALING | 42 | 0.6928073 | 2.67057 | 0.001148 | 0.00124794 | 5.25449E-05 | 11040 | tags=74%, list=22%, signal=58% |
| HALLMARK_ANGIOGENESIS | 36 | 0.680881921 | 2.57604 | 0.001144 | 0.00124794 | 5.25449E-05 | 12363 | tags=75%, list=24%, signal=57% |
| HALLMARK_BILE_ACID_METABOLISM | 112 | 0.573861392 | 2.54362 | 0.001016 | 0.00124794 | 5.25449E-05 | 13390 | tags=61%, list=27%, signal=45% |
| HALLMARK_HEDGEHOG_SIGNALING | 36 | 0.665837874 | 2.51912 | 0.001144 | 0.00124794 | 5.25449E-05 | 11040 | tags=56%, list=22%, signal=43% |
| HALLMARK_APICAL_SURFACE | 43 | 0.606202059 | 2.35731 | 0.00113 | 0.00124794 | 5.25449E-05 | 14010 | tags=60%, list=28%, signal=44% |
| HALLMARK_NOTCH_SIGNALING | 32 | 0.744163832 | 2.75143 | 0.001175 | 0.00125009 | 5.26355E-05 | 10601 | tags=84%, list=21%, signal=67% |
| HALLMARK_SPERMATOGENESIS | 132 | 0.345210263 | 1.54878 | 0.008081 | 0.00841751 | 0.000354421 | 12604 | tags=35%, list=25%, signal=26% |

**Supplementary Table 4 CMS of the 12 FF-FFPE-matched samples (AMC-FFPE CRC).**

| FF Sample Name | FFPE Sample Name | CMS (GSE33113) | CMS (FFPE) | CMS (FFPE-Rescuer) |
|---|---|---|---|---|
| GSM820058 | COL011 | CMS2 | CMS2 | CMS2 |
| GSM820061 | COL014 | CMS4 | CMS4 | CMS4 |
| GSM820067 | COL020 | CMS1 | CMS3 | CMS1 |
| GSM820085 | COL047 | CMS2 | CMS3 | CMS2 |
| GSM820089 | COL051_1 (S1) | CMS2 | CMS2 | CMS2 |
| GSM820089 | COL051_2 (rep) | CMS2 | CMS4 | CMS4 |
| GSM820121 | COL084 | CMS1 | CMS1 | CMS1 |
| GSM820125 | COL090 | CMS2 | CMS3 | CMS2 |
| GSM820127 | COL092 | CMS4 | CMS4 | CMS4 |
| GSM820133 | COL098_1 (S1) | CMS4 | CMS4 | CMS4 |
| GSM820133 | COL098_2 (rep) | CMS4 | CMS4 | CMS4 |
| GSM820135 | COL100 | CMS1 | CMS1 | CMS1 |

**Supplementary Table 5 Subtype of the 27 FFPE HGSOC samples (West China HGSOC).**

| FFPE Sample Name | Subtype (FFPE) | Subtype (FFPERescuer) |
|---|---|---|
| N116 | C1_immu | C4_mesc |
| N297 | C2_diff | C1_immu |
| N300 | C3_prof | C3_prof |
| N367 | C2_diff | C1_immu |
| N135 | C2_diff | C2_diff |
| N7_6583 | C2_diff | C2_diff |
| N8_10221 | C2_diff | C4_mesc |
| N8_10482 | C2_diff | C2_diff |
| NB10 | C2_diff | C2_diff |
| NB11 | C1_immu | C4_mesc |
| NB12 | C1_immu | C4_mesc |
| NB14 | C3_prof | C3_prof |
| NB18 | C2_diff | C3_prof |
| NB19 | C3_prof | C3_prof |
| N13 | C3_prof | C3_prof |
| NB21 | C3_prof | C3_prof |
| NB29 | C1_immu | C4_mesc |
| NB30 | C2_diff | C2_diff |
| NB35 | C2_diff | C2_diff |
| NB37 | C2_diff | C2_diff |
| NB39 | C2_diff | C1_immu |
| NB40 | C2_diff | C3_prof |
| N14 | C1_immu | C1_immu |
| N209 | C3_prof | C3_prof |
| N246 | C2_diff | C1_immu |
| N251 | C3_prof | C3_prof |
| N252 | C2_diff | C1_immu |

**Supplementary Table 6 A summary of public FFPE datasets.**

| Dataset | Sample size | Platform | Library preparation | Storage type | Accession |
|---|---|---|---|---|---|
| GSE139822 Bladder | 9 | Illumina HiSeq 3000 (Homo sapiens) | The Human FFPE RNA-seq kit (Total RNA) | FFPE | GSE139822 |
| GSE160693 Bladder | 52 | Illumina NextSeq 500 (Homo sapiens) | The Illumina RNA Access Library Prep Protocol (PolyA enrichment) | FFPE | GSE160693 |
| GSE146889 Colorectal | 76 | Illumina HiSeq 2500 (Homo sapiens) | TruSeq RNA exome library prep kit (Total RNA) | FFPE | GSE146889 |
| GSE152430 Colorectal | 49 | Illumina NextSeq 500 (Homo sapiens) | NEBNext Ultra II Directional Library Prep kit with NEBNext rRNA depletion module (rRNA depletion) | FFPE | GSE152430 |
| GSE86562 Colorectal | 70 | Illumina whole genome RNASeq (RNA-Acc); Illumina HiSeq 2000 | Illumina RNA-Access (PolyA enrichment) | FFPE | GSE86562 |
| GSE86564 Colorectal | 56 | Illumina HiSeq 2000 (Homo sapiens) | Illumina Total stranded RNA-rRNA-depletion (rRNA depletion) | FFPE | GSE86564 |
| GSE51124 Breast | 9 | Illumina HiSeq 2000 (Homo sapiens) | RiboZeroGold / ScriptSeq library preparation (rRNA depletion) | FFPE | GSE51124 |
| GSE113976 Breast | 14 | Illumina HiSeq 2500 (Homo sapiens) | Truseq RNA Sample preparation kit v2 (illumina) (PolyA enrichment); Ovation Human FFPE RNA-Seq Library (rRNA depletion) | FFPE | GSE113976 |
| GSE130397 Breast | 21 | Illumina HiSeq 2500 (Homo sapiens) | the TruSeq RNA Access Library Prep Kit(PolyA enrichment); the Ovation Human FFPE RNA-seq Library System (Total RNA) | FFPE | GSE130397 |
| GSE42948 Breast | 53 | Illumina Genome Analyzer IIx (Homo sapiens) | 3SEQ Sequencing library (PolyA enrichment) | FFPE | GSE42948 |

| | | | | | |
|---|---|---|---|---|---|
| GSE154041 GBM | 71 | Illumina NextSeq 500 (Homo sapiens) | the QuantSeq 3'mRNA-Seq Library Prep Kit-FWD (PolyA enrichment) | FFPE | GSE154041 |
| GSE54460 Prostate | 106 | Illumina HiSeq 2000 (Homo sapiens) | the TruSeq Kit (Illumina, Inc.) with the following modification. Instead of purifying poly-A RNA using poly-dT primer beads, we removed ribosomal RNA using the Ribominus Kit (rRNA depletion) | FFPE | GSE54460 |
| GSE89223 Prostate | 15 | Ion Torrent Proton (Homo sapiens) | Ion Total RNA-seq Kit v2 (Total RNA) | FFPE | GSE89223 |
| GSE148387 Conjunctival melanoma | 12 | Illumina NextSeq 500 (Homo sapiens) | Massive analysis of cDNA ends (MACE) library (PolyA enrichment) | FFPE | GSE148387 |
| GSE138866 Ovarian | 125 | HiSeq X Ten (Homo sapiens) | standard Illumina protocols (SMARTer® Stranded Total RNA-Seq Kit v2) (Total RNA) | FFPE | GSE138866 |
| GSE52248 Lung | 12 | Illumina Genome Analyzer IIx (Homo sapiens) | Ovation RNA-Seq FFPE (PolyA enrichment); Encore NGS Library System 1 kits (PolyA enrichment) | FFPE | GSE52248 |
| GSE143486 Lung | 24 | HiSeq X Ten (Homo sapiens) | NEBNext Ultra Directional RNA Library Prep Kit (Total RNA) | FFPE | GSE143486 |
| GSE147043 Gastric | 14 | Illumina NextSeq 500 (Homo sapiens) | QuantSeq 3′mRNA-Seq Library Prep Kit (PolyA enrichment) | FFPE | GSE147043 |
| Total | | | 788 | | |

**Supplementary Table 7 A summary of the validation datasets.**

| Dataset | Sample size | Cancer | Storage type | Library preparation | Platform | Resources |
|---|---|---|---|---|---|---|
| AMC-FFPE CRC | 12 | CRC | FFPE | Illumina Truseq RNA Exome Prep (Total RNA) | Illumina HiSeq 2500 | In-house |
| AMC-AJCCII-90 GSE33113 | 12 | CRC | Fresh Frozen | - | Affymetrix Human Genome U133 Plus 2.0 Array | Public |
| GSE86562 | 51 | CRC | FFPE | Illumina RNA-Access (Poly(A) enrichment) | Illumina whole genome RNASeq (RNA-Acc) Illumina HiSeq 2000 | Public |
| GSE86564 | 51 | CRC | FFPE | Illumina Total stranded RNA-rRNA-depletion (rRNA depletion) | Illumina HiSeq 2000 | Public |
| GSE86557 | 51 | CRC | Fresh Frozen | - | Affymetrix GeneChip | Public |
| West China HGSOC | 27 | HGSOC | FFPE | Poly(A) enrichment | Illumina HiSeq X Ten | In-house |

**Supplementary Table 8 A summary of public FF datasets used.**

| Dataset | Sample size | Storage type | Library preparation | Platform | Accession |
|---|---|---|---|---|---|
| The Cancer Genome Atlas (TCGA) | 9568 (tumor) | Fresh Frozen | - | - | - |
| GSE180440 | 145 | Fresh Frozen | Total RNA | Illumina HiSeq 2000 | GSE180440 |

**Supplementary Table 9 FF TCGA samples used for GSEA in simulation experiments.**

| Cancer type | Tumor size | Normal size |
|---|---|---|
| BLCA | 405 | 19 |
| BRCA | 1087 | 98 |
| COADREAD | 376 | 51 |
| KIPAN | 886 | 129 |
| LUAD | 514 | 58 |
| PRAD | 484 | 51 |
| Total | 3752 | 406 |
